# LZTR1 polymerization provokes cardiac pathology in recessive Noonan syndrome

**DOI:** 10.1101/2023.01.10.523203

**Authors:** Alexandra Viktoria Busley, Óscar Gutiérrez-Gutiérrez, Elke Hammer, Fabian Koitka, Amin Mirzaiebadizi, Martin Steinegger, Constantin Pape, Linda Böhmer, Henning Schroeder, Mandy Kleinsorge, Melanie Engler, Ion Cristian Cirstea, Lothar Gremer, Dieter Willbold, Janine Altmüller, Felix Marbach, Gerd Hasenfuss, Wolfram-Hubertus Zimmermann, Mohammad Reza Ahmadian, Bernd Wollnik, Lukas Cyganek

## Abstract

Noonan syndrome patients harboring causative variants in *LZTR1* are particularly at risk to develop severe and early-onset hypertrophic cardiomyopathy. However, the underling disease mechanisms of *LZTR1* missense variants driving the cardiac pathology are poorly understood. Hence, therapeutic options for Noonan syndrome patients are limited. In this study, we investigated the mechanistic consequences of a novel homozygous causative variant *LZTR1^L580P^*by using patient-specific and CRISPR/Cas9-corrected iPSC-cardiomyocytes. Molecular, cellular, and functional phenotyping in combination with *in silico* prediction of protein complexes uncovered a unique *LZTR1^L580P^*-specific disease mechanism provoking the cardiac hypertrophy. The homozygous variant was predicted to alter the binding affinity of the dimerization domains facilitating the formation of linear LZTR1 polymer chains. The altered polymerization resulted in dysfunction of the LZTR1-cullin 3 ubiquitin ligase complexes and subsequently, in accumulation of RAS GTPases, thereby provoking global pathological changes of the proteomic landscape ultimately leading to cellular hypertrophy. Furthermore, our data showed that cardiomyocyte-specific MRAS degradation is mediated by LZTR1 via the autophagosome, whereas RIT1 degradation is mediated by both LZTR1-dependent and LZTR1-independent proteasomal pathways. Importantly, uni-or biallelic genetic correction of the *LZTR1^L580P^* missense variant rescued the molecular and cellular disease-associated phenotype, providing proof-of-concept for CRISPR-based gene therapies.

## Introduction

Noonan syndrome (NS) is a multi-systemic developmental disorder with a broad spectrum of symptoms and varying degrees of disease severity. Common clinical symptoms range from intellectual disability, facial dysmorphisms, webbed neck, skeletal deformities, short stature, and in many cases congenital heart disease.^1^ With a prevalence of approximately 1 in 1,000 – 2,500 live births, NS is considered the most common monogenic disease associated with congenital heart defects and early-onset hypertrophic cardiomyopathy (HCM).^2^ Young NS patients diagnosed with HCM are more prone to develop heart failure accompanied by a poor late survival in contrast to patients suffering from non-syndromic HCM.^3,4^ Like other phenotypically overlapping syndromes classified as RASopathies, NS is caused by variants in RAS-mitogen-activated protein kinase (MAPK)-associated genes, all typically leading to an increase in signaling transduction.^5^ Within the RASopathy spectrum, patients harboring causative gene variants in *RAF1*, *HRAS*, *RIT1* and *LZTR1* are particularly at risk to develop severe and early-onset HCM.^6,7^

Recent studies by others and our group have revealed the functional role of LZTR1 within the RAS-MAPK signaling cascade as a negative regulator of signaling activity. *LZTR1* encodes an adapter protein of the cullin 3 ubiquitin ligase complex by selectively targeting RAS proteins as substrates for degradation. *LZTR1* deficiency – caused by truncating or missense variants – results in an accumulation of the RAS protein pool and, as a consequence, in RAS-MAPK signaling hyperactivity.^8–10^ Whereas dominant *LZTR1* variants generally cluster in the Kelch motif perturbing RAS binding to the ubiquitination complex,^11^ the mechanistic consequences of recessive *LZTR1* missense variants, which are distributed over the entire protein, are not understood.

Human induced pluripotent stem cell-derived cardiomyocytes (iPSC-CMs) generated from patients with inherited forms of cardiomyopathies offer a unique platform to study the disease mechanisms in physiologically relevant cells and tissues.^12,13^ A few RASopathy-linked iPSC-CM models had been described, including for variants in *PTPN11*, *RAF1*, *BRAF*, and *MRAS*.^14–17^ In line, we had recently added novel information as to the role of *LZTR1*-truncating variants in NS pathophysiology.^10,18^ In the present study, we aimed to investigate the molecular, cellular, and functional consequences of a specific recessive missense variant *LZTR1^L580P^* by utilizing patient-derived and CRISPR-corrected iPSC-CMs. We could show that *LZTR1^L580P^*in homozygous state results in aberrant polymerization causing LZTR1 dysfunction, marked increase of RAS GTPase levels, and cellular hypertrophy. Further, uni-and bi-allelic genetic correction of the missense variant by CRISPR/Cas9 technology rescued the cellular phenotype, indicating that correction of one allele is sufficient to restore the cardiac pathophysiology, thereby providing proof-of-concept for future personalized CRISPR-based gene therapies.

## Results

### LZTR1^L580P^ is causative for recessive Noonan syndrome

A 17-year old male patient with HCM, stress-induced cardiac arrhythmias, pectus excavatum and facial anomalies was referred to our clinic, and based on the combination of symptoms, the clinical diagnosis of Noonan syndrome was made (Figure 1A, and Table S1 in the supplement). The patient was born to a consanguineous couple and both parents showed neither apparent clinical symptoms nor distinctive NS-specific features. Whole exome sequencing followed by detailed variant analysis detected one highly suspicious homozygous variant in *LZTR1*. Both parents were heterozygous carriers and the variant was not present in any current database of human genetic variations including the >250,000 alleles of gnomAD database. The homozygous missense variant, c.1739T>C, was located in exon 15 of the *LZTR1* gene and leads to the substitution of an evolutionary conserved leucine at the amino acid position 580 by proline (p.Leu580Pro, p.L580P). The variant was predicted as likely pathogenic by computational predictions.

**Figure 1:**
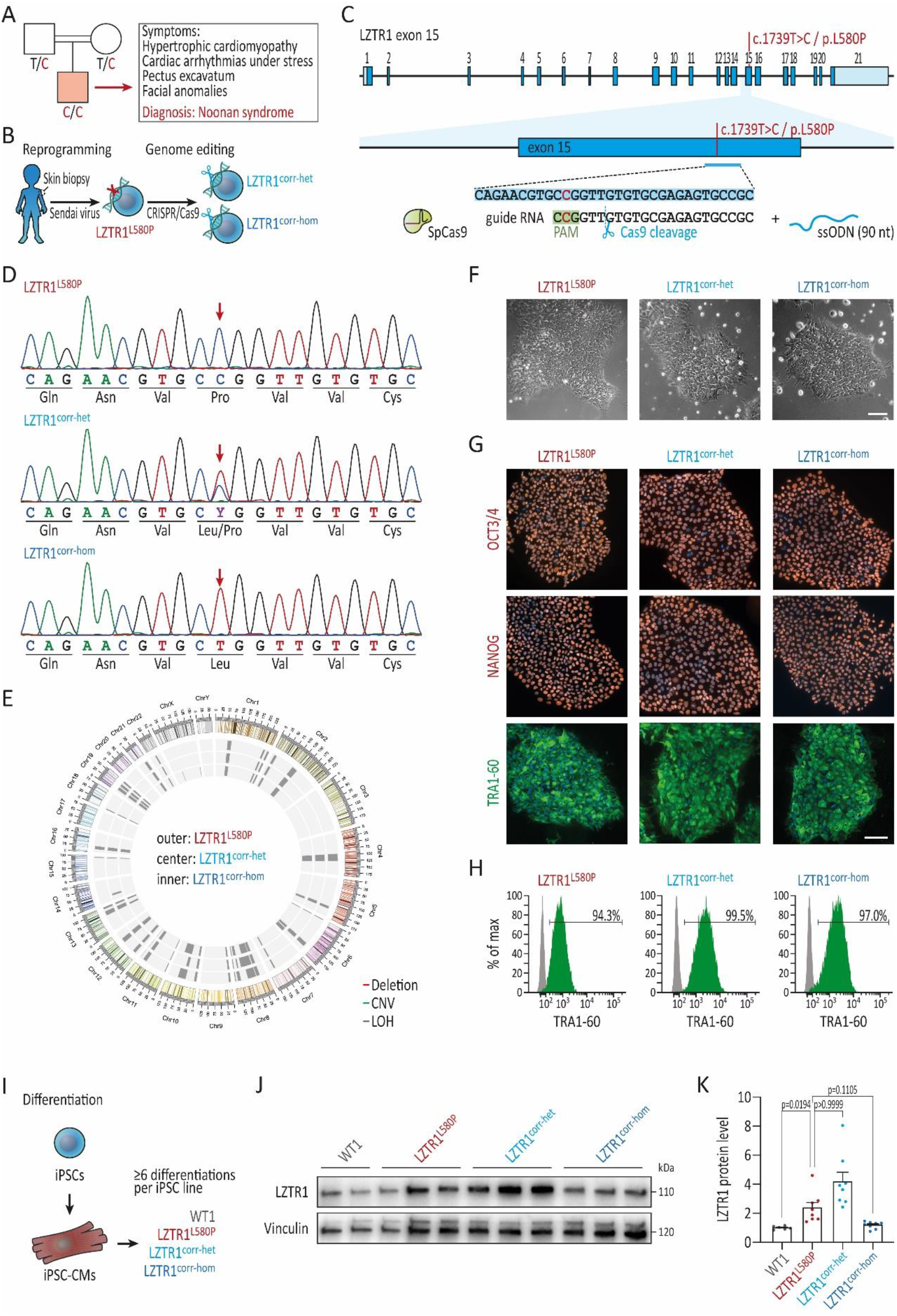
Generation of patient-specific and CRISPR-corrected iPSCs for disease modeling of recessive NS. **(A)** Pedigree of the consanguineous family with healthy parents and the son affected by recessive NS harboring the *LZTR1* variant (c.1739T>C/p.L580P) in homozygosity. **(B)** Generation of patient-specific iPSCs by reprogramming of patient’s skin fibroblasts via integration-free Sendai virus and genetic correction of the missense variant by CRISPR/Cas9. **(C)** Depiction of the genome editing approach for correction of the missense variant in *LZTR1* exon 15 by CRISPR/Cas9 and single-stranded oligonucleotide (ssODN) for homology-directed repair. **(D)** Sanger sequencing of the patient-derived iPSCs (LZTR1^L580P^) with the *LZTR1* missense variant in homozygosity and the CRISPR/Cas9-edited heterozygous corrected (LZTR1^corr-het^) and homozygous corrected (LZTR1^corr-hom^) iPSCs. **(E)** Molecular karyotyping using a genome-wide microarray demonstrated a high percentage of loss of heterozygosity (LOH) because of consanguinity as well as chromosomal stability of iPSCs after genome editing. **(F)** Patient-specific and CRISPR-corrected iPSCs showed a typical human stem cell-like morphology; scale bar: 100 µm. **(G)** Expression of key pluripotency markers OCT3/4, NANOG, and TRA-1-60 in the generated iPSC lines was assessed by immunocytochemistry; nuclei were counter-stained with Hoechst 33342 (blue); scale bar: 100 μm. **(H)** Flow cytometry analysis of pluripotency marker TRA-1-60 revealed homogeneous populations of pluripotent cells in generated iPSC lines. Gray peaks represent the negative controls. **(I)** Differentiation of WT, patient-specific and CRISPR-corrected iPSCs into iPSC-CMs. **(J)** Representative blot of endogenous LZTR1 levels in WT, patient’s, and CRISPR-corrected iPSC-CMs at day 60 of differentiation, assessed by Western blot; Vinculin served as loading control; n=3 individual differentiations per iPSC line. **(K)** Quantitative analysis of Western blots for LZTR1; data were normalized to total protein and to the corresponding WT samples on each membrane; n=6-8 independent differentiations per iPSC line. Data were analyzed by nonparametric Kruskal-Wallis test with Dunn correction and are presented as mean ± SEM (K).

To elucidate the molecular and functional consequences of the *LZTR1^L580P^*missense variant, we generated iPSCs from the patient’s skin fibroblasts using integration-free reprograming methods and subsequently utilized CRISPR/Cas9 genome editing to engineer gene variant-corrected iPSC lines (Figure 1B). For genetic correction of the patient-specific iPSCs, the CRISPR guide RNA was designed to specifically target the mutated sequence in exon 15 of the *LZTR1* gene. Further, the ribonucleoprotein-based CRISPR/Cas9 complex was combined with a single-stranded oligonucleotide serving as template for homology-directed repair (Figure 1C). Upon transfection, cells were singularized and individual clones were screened for successful editing to identify heterozygous corrected as well as homozygous corrected iPSC clones, LZTR1^corr-het^ and LZTR1^corr-hom^, respectively (Figure 1D). Molecular karyotyping of the edited iPSC clones confirmed chromosomal stability after genome editing and passaging (Figure 1E). As expected for individuals born to consanguineous parents, patient-specific as well as CRISPR-corrected iPSCs demonstrated a noticeable reduction of the overall heterozygosity using SNP-based genome-wide arrays, with around 30% of segments of the genome being assigned to regions of heterozygosity. Further, sequencing revealed no obvious off-target modifications by genome editing (Figure S1 in the supplement). Subsequently, patient-derived and CRISPR-corrected iPSCs were verified for pluripotency (Figure 1F-H). In addition to the patient-derived iPSC lines, iPSC lines from two unrelated healthy male donors, namely WT1 and WT11, were used as wild type (WT) controls in this study.

At first, we aimed to determine whether the LZTR1^L580P^ protein remains stably expressed or is rapidly degraded after protein translation. LZTR1 proteins were robustly detected by Western blot in differentiated iPSC-CMs (Figure 1I-K). Interestingly, significantly higher LZTR1 protein levels were present in the patient-specific and the heterozygous corrected iPSC-CMs compared to WT and homozygous corrected cultures, suggesting an accumulation of the mutant LZTR1^L580P^ proteins in the cells.

### Homozygous LZTR1^L580P^ causes accumulation of RAS GTPases

To investigate the impact of the identified homozygous *LZTR1^L580P^*missense variant on the molecular mechanisms contributing to left ventricular hypertrophy, patient-specific, heterozygous and homozygous corrected, as well as two individual WT iPSC lines were differentiated into functional ventricular-like iPSC-CMs in feeder-free culture conditions,^19^ and on day 60 of differentiation subjected to unbiased proteome analyses (Figure 2A). We identified more than 4,700 proteins in the samples from the individual groups. All samples showed a comparably high abundance of prominent cardiac markers including myosin heavy chain β (*MHY7*), cardiac troponin T (*TNNT2*), α-actinin (*ACTN2*), titin (*TTN*), and ventricular-specific myosin light chain 2 (*MYL2*), indicating equal cardiomyocyte content in the different cultures (Figure 2B). By comparing the proteome profiles of LZTR1^L580P^ and WT iPSC-CMs, we identified enhanced abundance of the RAS family members muscle RAS oncogene homolog (MRAS) and RIT1 in the patient’s iPSC-CMs (Figure 2C). This finding is in agreement with our previous observation in *LZTR1*-truncating variant carriers^10^ and confirms the pivotal role of LZTR1 in targeting various RAS GTPases for LZTR1-cullin 3 ubiquitin ligase complex-mediated ubiquitination, and degradation.^8,9^ Further, it highlights that *LZTR1^L580P^*results in protein loss-of-function, causing an accumulation of RAS proteins in the cells, providing molecular evidence for the causative nature of the missense variant. Strikingly, protein levels of the different RAS GTPases were normalized in both the heterozygous as well as the homozygous corrected iPSC-CMs, confirming that only one functional *LZTR1* allele is sufficient to regulate the protein pool of RAS GTPases in cardiomyocytes (Figure 2D-E). As anticipated, transcriptome analyses showed similar mRNA expression levels of the different RAS GTPases in the patient’s and CRISPR-corrected iPSC lines, indicating a post-translational cause for the higher abundance of RAS proteins in LZTR1^L580P^ cultures (Figure S2 in the supplement). In contrast, the significantly elevated protein levels of the protein quality control-associated heat shock-related 70 kDa protein 2 (*HSPA2*) in the patient’s cells in comparison to the WT and CRISPR-corrected cells were related to upregulation of gene expression, suggesting that HSPA2 is not directly targeted by LZTR1 for degradation.

**Figure 2:**
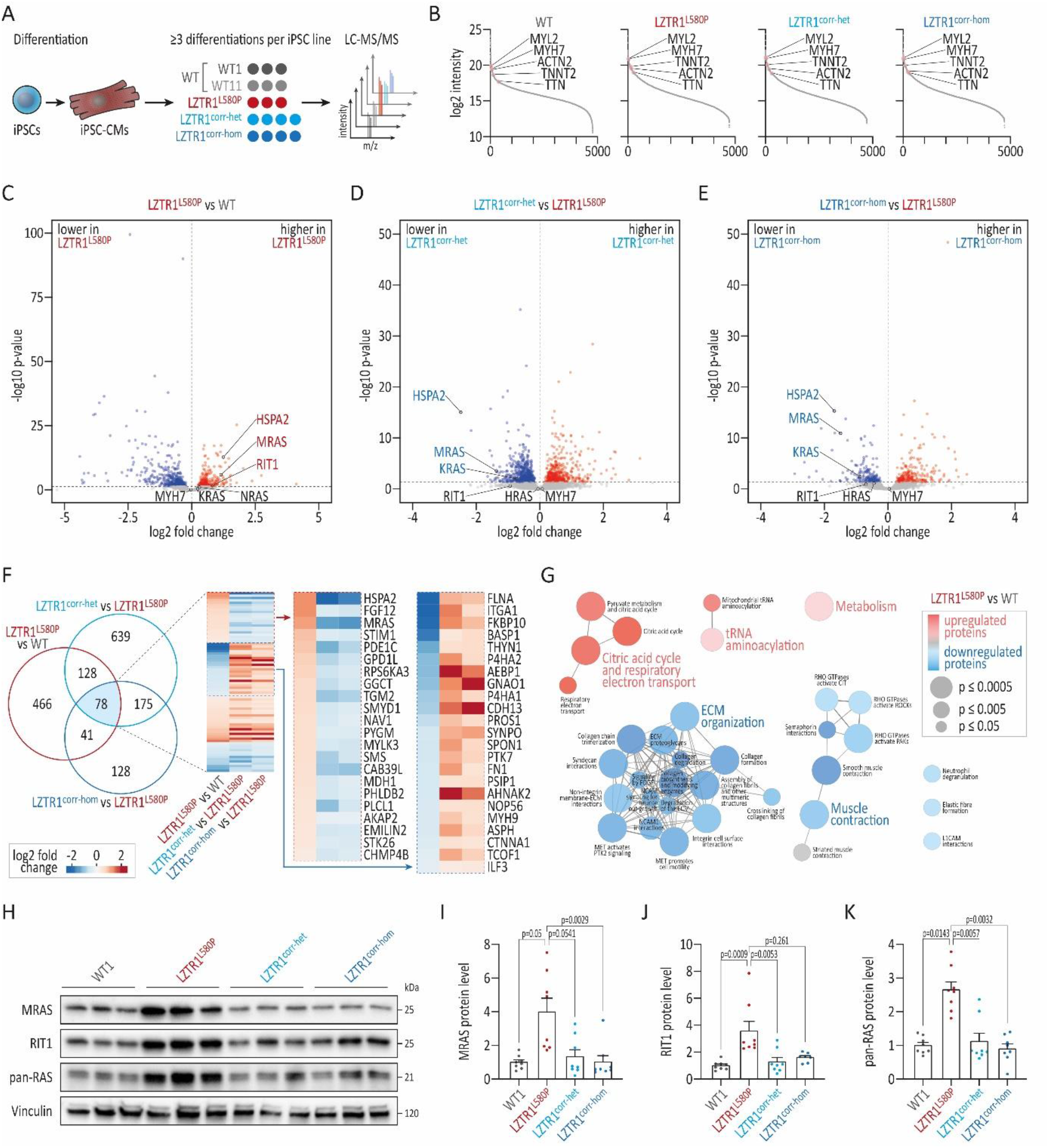
Homozygous *LZTR1^L580P^*causes accumulation of RAS GTPases. **(A)** Depiction of the experimental design: two individual WT, the patient-specific, and the two CRISPR-corrected iPSC lines were differentiated into ventricular iPSC-CMs and analyzed by quantitative global proteomics via LC-MS/MS at day 60 of differentiation; n=3-4 individual differentiations per iPSC line. **(B)** Over 4,700 proteins were present in the individual proteomic samples, all showing comparable high abundance of cardiac markers myosin heavy chain β (*MHY7*), cardiac troponin T (*TNNT2*), α-actinin (*ACTN2*), titin (*TTN*), and ventricular-specific MLC2V (*MYL2*). **(C-E)** Volcano plots representing relative protein abundances comparing patient’s versus WT iPSC-CMs (C; LZTR1^L580P^ vs WT), heterozygous corrected versus non-corrected iPSC-CMs (D; LZTR1^corr-het^ vs LZTR1^L580P^), and homozygous corrected versus non-corrected iPSC-CMs (E; LZTR1^corr-hom^ vs LZTR1^L580P^) revealed high abundance of RAS GTPases in patient samples. **(F)** Comparison of differentially abundant proteins between the three datasets revealed an overlap of 78 proteins, many of which showed opposite abundance in patient’s versus CRISPR-corrected iPSC-CMs. **(G)** Reactome pathway enrichment analysis of differentially abundant proteins in LZTR1^L580P^ vs WT displayed dysregulation of cardiac-related pathways and biological processes. **(H)** Representative blots of RAS GTPase levels in WT, patient’s, and CRISPR-corrected iPSC-CMs at day 60 of differentiation, assessed by Western blot; Vinculin served as loading control; n=3 individual differentiations per iPSC line. **(I-K)** Quantitative analysis of Western blots for MRAS (I), RIT1 (J), and pan-RAS recognizing HRAS, KRAS, and NRAS (K); data were normalized to total protein and to the corresponding WT samples on each membrane; n=8 independent differentiations per iPSC line. Data were analyzed by nonparametric Kruskal-Wallis test with Dunn correction and are presented as mean ± SEM (I-K).

To assess the correlation of the different proteomic profiles with respect to the disease-specific proteome signatures upon *LZTR1* deficiency, we performed a comparison analysis of the data sets from (1) LZTR1^L580P^ versus WT, (2) LZTR1^corr-het^ versus LZTR1^L580P^, and (3) LZTR1^corr-hom^ versus LZTR1^L580P^. We found 78 proteins being differentially regulated in all three data sets (Figure 2F). Here, a profound subset of proteins of the overlapping profile that was significantly higher abundant in the patient’s cells, such as the MAPK-activated protein kinase RPS6KA3, was normalized after heterozygous and homozygous CRISPR-correction of the pathological *LZTR1* variant. Vice versa, numerous downregulated proteins in the patient samples were found to be elevated in the gene-edited iPSC-CMs. Further, we performed a Reactome pathway enrichment analysis to uncover dysregulated pathways and/or biological processes associated to *LZTR1^L580P^*. The analysis indicated that differentially abundant proteins in patient-derived samples were enriched in critical cardiac-related biological processes, such as muscle contraction and extracellular matrix organization, as well as in cellular routes associated to metabolism (Figure 2G). In agreement with the proteomic data, Western blot analysis confirmed the strong accumulation of MRAS, RIT1, and the classical RAS GTPases (HRAS, KRAS, and NRAS; detected by pan-RAS) in the LZTR1^L580P^ cultures, and further confirmed a normalization of RAS levels in the CRISPR-corrected isogenic iPSC-CMs to WT control levels (Figure 2H-K).

Collectively, these data demonstrate that the missense variant *LZTR1^L580P^* in homozygosity resulted in protein loss-of-function causing an accumulation of RAS GTPases as the critical underlying disease mechanism in cardiomyocytes from the NS patient. In line, correction of the homozygous missense variant on at least one allele normalized the molecular pathology.

### Homozygous LZTR1^L580P^ retains a residual protein function

To explore the impact of RAS GTPase accumulation on RAS-MAPK signaling activity, we used an ERK kinase translocation reporter (ERK-KTR) to measure ERK signaling dynamics in live cells.^20^ Patient-specific, heterozygous and homozygous corrected, and WT iPSC-CMs were efficiently transduced with the ERK-KTR lentivirus, and the activity of ERK was analyzed at day 60 of differentiation by measuring the ratio of cytosolic (corresponding to active ERK) to nuclear (corresponding to inactive ERK) fluorescent signals (Figure 3A,B). The specificity of the ERK biosensor was confirmed by a selective response to MEK inhibition, whereas no change in ERK biosensor activity was observed when cells were treated with an inhibitor of the JNK pathway (Figure S3 in the supplement). Biosensor-transduced iPSC-CM cultures were treated with the MEK inhibitor trametinib (MEKi) or with DMSO for 60 minutes, before stimulation with fetal bovine serum for another 60 minutes, and imaged every 10 minutes (Figure 3C, and Figure S3 in the supplement). Under basal conditions, an equally low level of ERK activity was observed across all iPSC lines (Figure 3D). As expected, a strong increase in ERK activity was detected upon stimulation of the cells, while MEK inhibition was effective in normalizing ERK signaling activity (Figure 3D). The results of the imaging-based approach were confirmed by Western blot analysis of uncorrected and CRISPR-corrected iPSC-CMs (Figure 3E).

**Figure 3:**
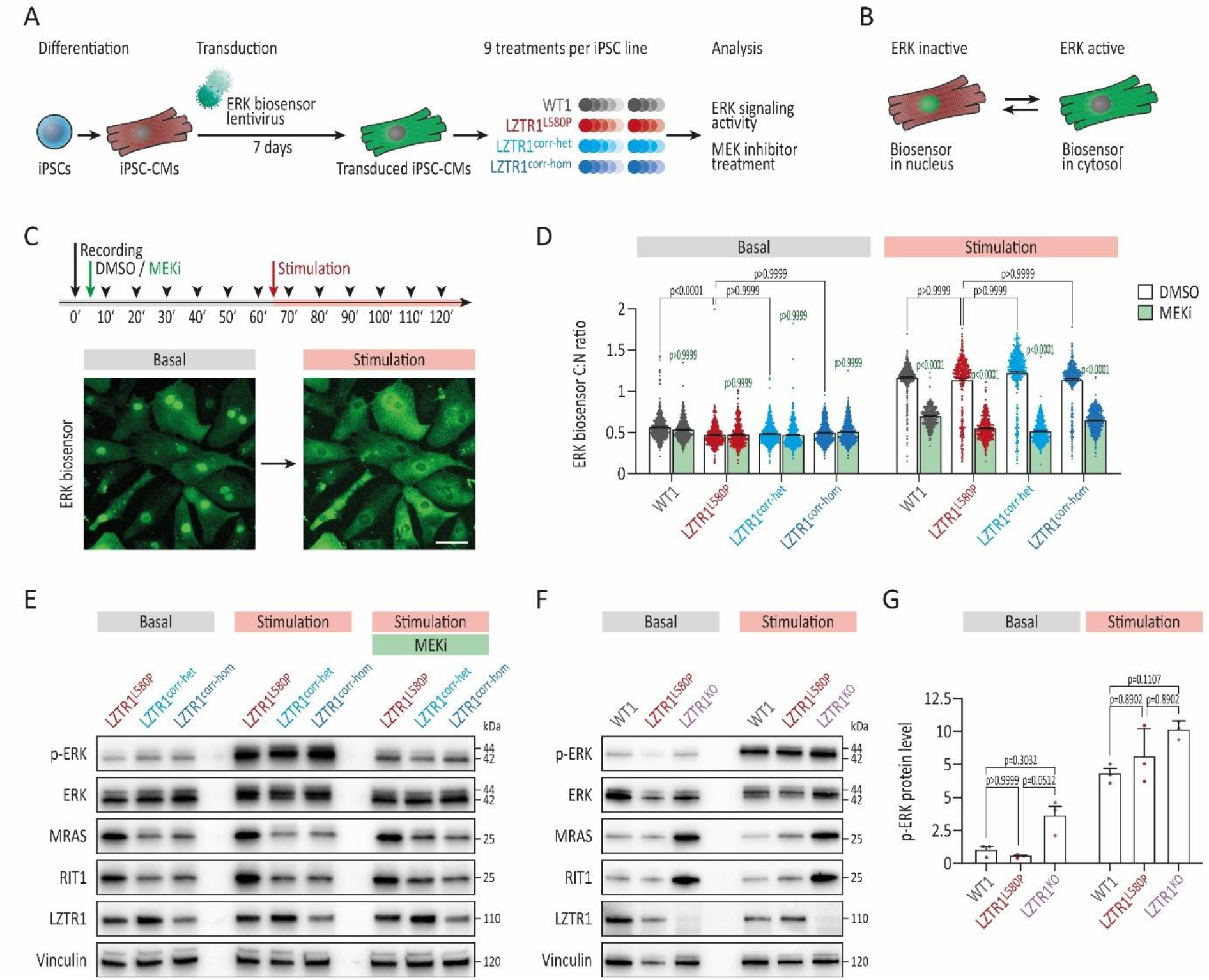
Homozygous *LZTR1^L580P^* retains a residual protein function. **(A)** Depiction of the experimental design: the WT, the patient-specific, and the two CRISPR-corrected iPSC lines were differentiated into ventricular iPSC-CMs and transduced around day 50 of differentiation with lentivirus containing an ERK kinase translocation reporter (ERK-KTR) to measure ERK signaling dynamics in real time. **(B)** ERK activity was analyzed by measuring the ratio of cytosolic (corresponding to active ERK) to nuclear (corresponding to inactive ERK) fluorescent signals. **(C)** Biosensor-transduced iPSC-CMs were treated with MEK inhibitor trametinib (MEKi) or with DMSO for 60 minutes, before stimulation with serum for another 60 minutes, and imaged every 10 minutes. **(D)** Quantitative analysis of ERK biosensor cytosol/nucleus (C:N) ratio under basal conditions (60 minutes after MEKi/DMSO treatment) and 20 minutes after stimulation; n=2 independent differentiations per iPSC line with n=4-5 individual wells per condition. **(E)** Representative blots of p-ERK, ERK, MRAS, RIT1, and LZTR1 levels in patient’s and CRISPR-corrected iPSC-CMs at day 60 of differentiation under basal conditions and 30 minutes after stimulation with and without pre-treatment with MEKi, assessed by Western blot; Vinculin served as loading control. **(F)** Representative blots of p-ERK, ERK, MRAS, RIT1, and LZTR1 levels in WT, patient’s and LZTR1^KO^ iPSC-CMs at day 60 of differentiation under basal conditions and 30 minutes after stimulation, assessed by Western blot; Vinculin served as loading control. **(G)** Quantitative analysis of Western blots for p-ERK protein levels; data were normalized to total protein and to the corresponding WT samples on each membrane; n=3 independent differentiations per iPSC line. Data were analyzed by nonparametric Kruskal-Wallis test with Dunn correction and are presented as mean ± SEM (D, G).

Since we did not observe increased ERK activity attributed to the homozygous *LZTR1^L580P^* missense variant, we compared the patient-specific LZTR1^L580P^ cells with another patient line harboring biallelic *LZTR1* variants causing complete loss of LZTR1 protein expression (LZTR1^KO^), which we reported in our previous study.^10^ Here, higher levels of phosphorylated ERK were observed in the LZTR1^KO^ cultures under basal conditions and after stimulation (Figure 3F,G). Interestingly, LZTR1^KO^ iPSC-CMs exhibited a substantially higher accumulation of RAS GTPases compared to LZTR1^L580P^ cells, implying a partial residual function of LZTR1^L580P^ ubiquitin ligase complexes.

### Homozygous LZTR1^L580P^ provokes cardiomyocyte hypertrophy

To elucidate the consequences of dysregulated RAS-MAPK signaling on the cellular characteristics of cardiomyocytes, we investigated the sarcomere homogeneity, the overall myofibril organization, and the cell size of the patient-derived iPSC-CMs, the CRISPR-corrected cells, as well as the WT controls at day 60 of differentiation (Figure 4A). Immunocytochemical staining of cardiac subtype-specific proteins revealed that all iPSC lines exhibit a well-organized sarcomeric organization with a pronounced striated expression of α-actinin and ventricular-specific MLC2V (Figure 4B). In order to analyze the homogeneity of sarcomeres in detail, we measured the distances between the sarcomeric Z-disks along individual myofibrils (Figure 4C). In agreement with the sarcomere length previously observed in neonatal and adult human hearts,^21^ *LZTR1*-deficient as well as corrected and WT cells revealed a typical sarcomere length ranging from 1.7 to 2.2 µm with an average of approximately 1.9 µm across all iPSC lines (Figure 4D). As sarcomeric disarray has been frequently reported in other iPSC-CM models of both NS-associated and non-syndromic HCM,^15,22^ we examined the myofibril organization in the individual iPSC-CMs stained for α-actinin by Fast Fourier Transform. The quantitative analysis did neither reveal any decrease of sarcomere regularity nor any pathological myofibril organization in LZTR1^L580P^ cultures (Figure 4E). On the contrary, LZTR1^L580P^ and CRISPR-corrected iPSC-CMs even demonstrated a slightly higher myofibril regularity compared to unrelated controls, indicating that the pathological gene variant has no severe impact on sarcomere structures.

**Figure 4:**
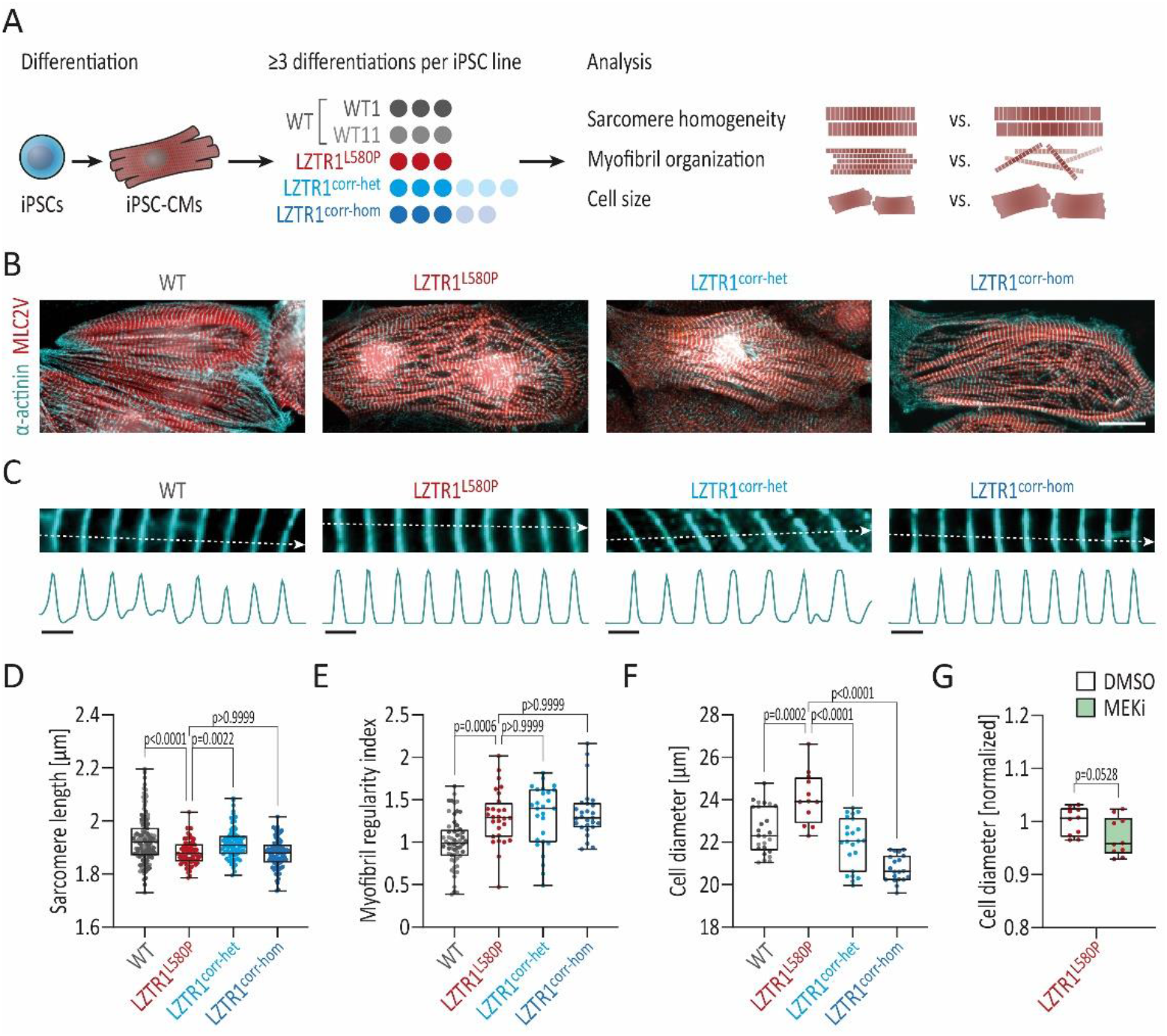
Homozygous *LZTR1^L580P^*provokes cardiomyocyte hypertrophy. **(A)** Depiction of the experimental design: two individual WT, the patient-specific, and the two CRISPR-corrected iPSC lines were differentiated into ventricular iPSC-CMs and analyzed for sarcomere length, myofibril organization, and cell size at day 60 of differentiation. **(B)** Representative images of iPSC-CMs stained for α-actinin and ventricular-specific MLC2V indicated a regular and well-organized sarcomeric assembly across all iPSC lines; scale bar: 20 µm. **(C)** Analysis of the mean sarcomere length per cell was based on measurement of multiple α-actinin-stained individual myofibrils; representative myofibrils and corresponding intensity plots are shown; scale bar: 2 µm. **(D)** Quantitative analysis displayed a typical sarcomere length in iPSC-CMs ranging from 1.7 to 2.2 µm across all iPSC lines; n=75-135 cells from 3 individual differentiations per iPSC line. **(E)** Quantitative analysis of the myofibril organization in α-actinin-stained iPSC-CMs, assessed by Fast Fourier Transform algorithm, demonstrated a high myofibril regularity across all iPSC lines; data were normalized to WT; n=27-58 images from 3 individual differentiations per iPSC line. **(F)** Quantitative analysis of the cell diameter in suspension in singularized iPSC-CMs, assessed by CASY cell counter, revealed a hypertrophic cell diameter in patient’s cells, compared with WT and CRISPR-corrected iPSC-CMs; n=12-25 samples from 3-6 individual differentiations per iPSC line. **(G)** Quantitative analysis of the cell diameter in suspension in singularized patient-specific iPSC-CMs that were treated with MEK inhibitor trametinib (MEKi) or with DMSO for 5 days, assessed by CASY cell counter; n=3 independent differentiations with n=3-4 individual wells per condition. Data were analyzed by nonparametric Kruskal-Wallis test with Dunn correction (D-F) or unpaired t test (G) and are presented as mean ± SEM.

As cardiomyocyte hypertrophy is a major hallmark of HCM, we further investigated the medium cell size of iPSC-CMs from all cell lines by utilizing our previously established assay to determine the cell size of iPSC-CMs in suspension.^10^ Here, the patient’s iPSC-CMs displayed a significant cellular enlargement compared to WT iPSC-CMs (Figure 4F). Strikingly, the hypertrophic phenotype was normalized in the CRISPR-corrected cells from both the LZTR1^corr-het^ and the LZTR1^corr-hom^ isogenic cultures. Moreover, and in line with the molecular observations, heterozygous correction of the pathological variant was sufficient to significantly reduce cellular hypertrophy. Additionally, we assessed whether treatment with the MEK inhibitor trametinib for 5 days could reverse the cellular hypertrophy in the patient-specific iPSC-CMs (Figure 4G). No significant reduction in cell size was observed in MEKi-treated cells compared to DMSO-treated cells, suggesting that normalization of RAS-MAPK signaling activity is unable to alleviate the cellular pathology in the short term.

In summary, the patient’s iPSC-CMs harboring the homozygous missense variant *LZTR1^L580P^*recapitulated the cardiomyocyte hypertrophy *in vitro*. Importantly, CRISPR-correction of the pathological variant was able to normalize the hypertrophic phenotype.

### Homozygous LZTR1^L580P^ does not compromise contractile function

NS-associated HCM as well as inherited forms of non-syndromic HCM are frequently associated with contractile dysfunction and these patients are at risk for developing arrhythmias.^23,24^ Hence, we generated engineered heart muscles (EHMs) from diseased, CRISPR-corrected, and WT iPSC-CMs enabling us to investigate the functional characteristics in a three-dimensional environment closer resembling the native conditions of the human heart muscle (Figure 5A).^25,26^ Microscopically, all iPSC lines formed homogenous cardiac tissues without showing apparent cell line-dependent differences after six weeks of cultivation and maturation (Figure 5B). Optical measurements were performed to study beating rate, force of contraction, and contraction kinetics in spontaneously contracting EHMs (Figure 5C). In comparison to WT EHMs, an increased spontaneous beat frequency was detected in the LZTR1^L580P^ EHMs (Figure 5D). The beat rate acceleration was gradually normalized in the heterozygous and homozygous corrected variants. Low beat-to-beat variability indicated that the LZTR1 mutant tissues do not provoke arrhythmia (Figure 5E). No significant differences in force of contraction were identified (Figure 5F). In accordance with higher beat frequencies, an acceleration of contraction and relaxation kinetics were observed in LZTR1^L580P^-, LZTR1^corr-het^-and the LZTR1^corr-hom^-derived EHMs (Figure 5G,H). However, since the altered kinetics were noticed in both diseased and CRISPR-corrected tissues, this rather suggested a mutation-independent effect. In addition, we examined the contractile properties of 2D cardiac monolayer cultures by video analysis and did not observe any significant differences between WT, patient-specific, CRISPR-corrected, and LZTR1^KO^ iPSC lines (Figure S4 in the supplement). Taken together, this functional data indicates that the missense variant *LZTR1^L580P^*does not significantly impact the contractile function and rhythmogenesis of cardiomyocytes.

**Figure 5:**
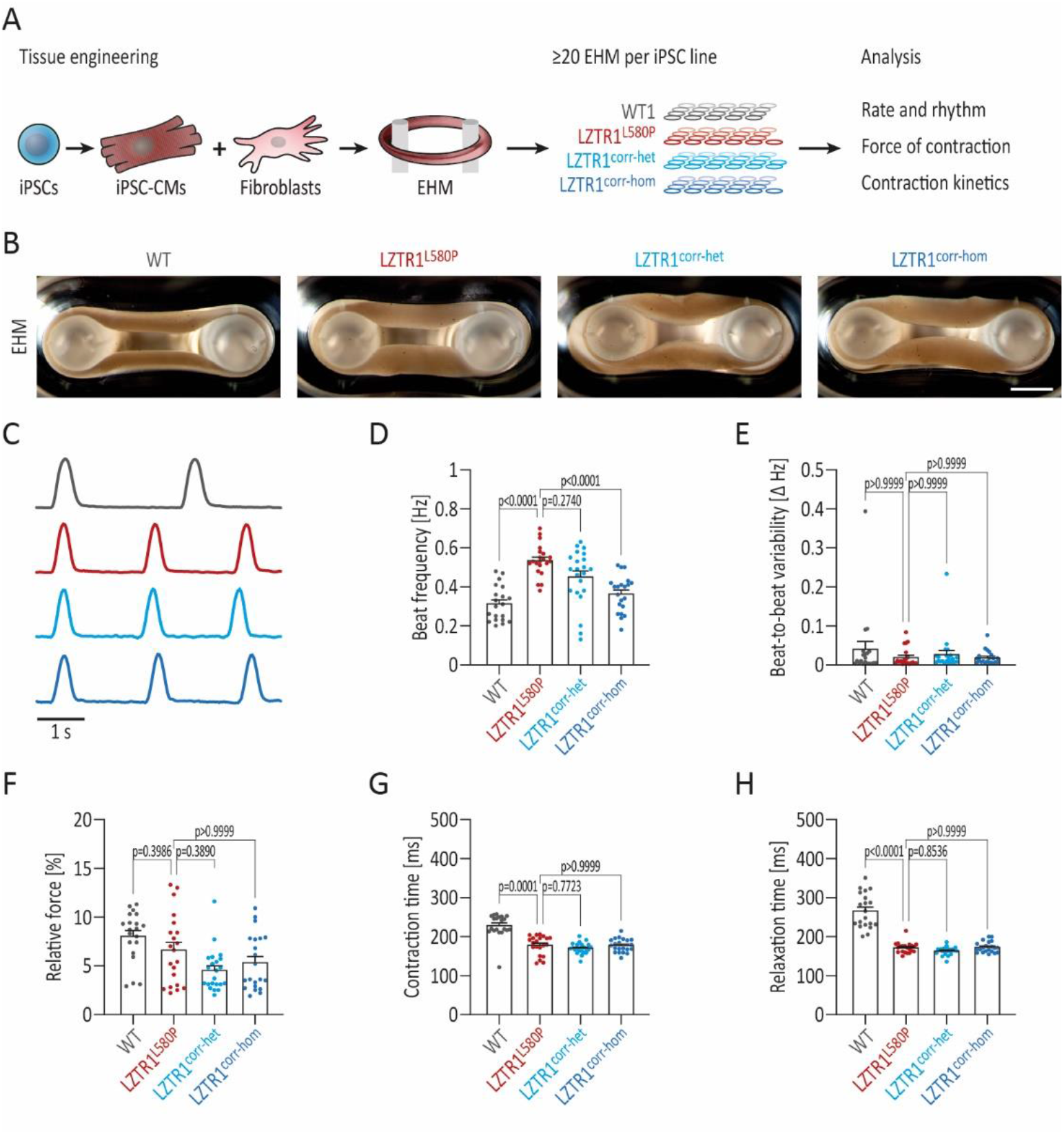
Homozygous *LZTR1^L580P^*does not compromise contractile function. **(A)** Depiction of the experimental design: the WT, the patient-specific, and the two CRISPR-corrected iPSC lines were differentiated into ventricular iPSC-CMs and casted at day 30 of differentiation together with fibroblasts in a collagen matrix for generation of EHMs. Tissues were analyzed for rhythmogenicity and contractile parameters by optical recordings at 5-6 weeks post-casting; n=20-22 EHMs from 3 individual differentiations per iPSC line. **(B)** Representative microscopic images of generated EHMs 6 weeks post-casting showing comparable tissue morphologies; scale bar: 1 mm. **(C)** Exemplary contraction traces from optical recordings of EHMs 6 weeks post-casting; peak amplitudes were normalized. **(D)** Quantitative analysis of the beating frequency of spontaneously contracting EHMs displayed minor differences in patient-derived tissues. **(E)** Quantitative measurement of the beat-to-beat variability of spontaneously contracting EHMs showed equal beating regularities across all tissues. **(F)** Quantitative analysis of the force of contraction, assessed by measuring the relative deflection of flexible poles, identified no significant differences across all iPSC lines. **(G-H)** Quantitative analysis of the contraction kinetics revealed longer contraction times (G) and relaxation times (H) in WT compared to patient’s and CRISPR-corrected EHMs. Data were analyzed by nonparametric Kruskal-Wallis test with Dunn correction and are presented as mean ± SEM (D-H).

### Homozygous LZTR1^L580P^ induces polymerization of LZTR1-cullin 3 ubiquitin ligase complexes

Considering the severe consequence of *LZTR1^L580P^* on the molecular and cellular pathophysiology in cardiomyocytes, we aimed to determine the specific effect of this variant on protein structure, complex formation, as well as its subcellular localization. We were not able to visualize endogenous LZTR1 in our cell model by immunocytochemistry, neither by testing of multiple commercial antibodies nor by N-terminal or C-terminal genetic tagging of the *LZTR1* gene locus. In order to circumvent these obstacles, we established ectopic expression of tagged *LZTR1* in WT iPSC-CMs at around day 60 of differentiation by lipofectamine-based plasmid transfection (Figure 6A). Besides *LZTR1^WT^* and *LZTR1^L580P^*, we screened the NS patient database^27^ for additional missense variants classified as likely pathogenic or variant of uncertain significance and located in close proximity to *LZTR1^L580P^* (within the BACK1 domain), and included these in our screening panel (Figure 6B). Of note, except for *LZTR1^L580P^* and *LZTR1^E563Q^*,^7^ none of the other variants had been reported to be present in homozygosity in *LZTR1*-associated NS. In addition, we also included a truncating variant *LZTR1^ΔBTB2-BACK2^*, lacking the entire BTB2-BACK2 domain, that mimicked the genotype of the two siblings described in our previous study.^10^

**Figure 6:**
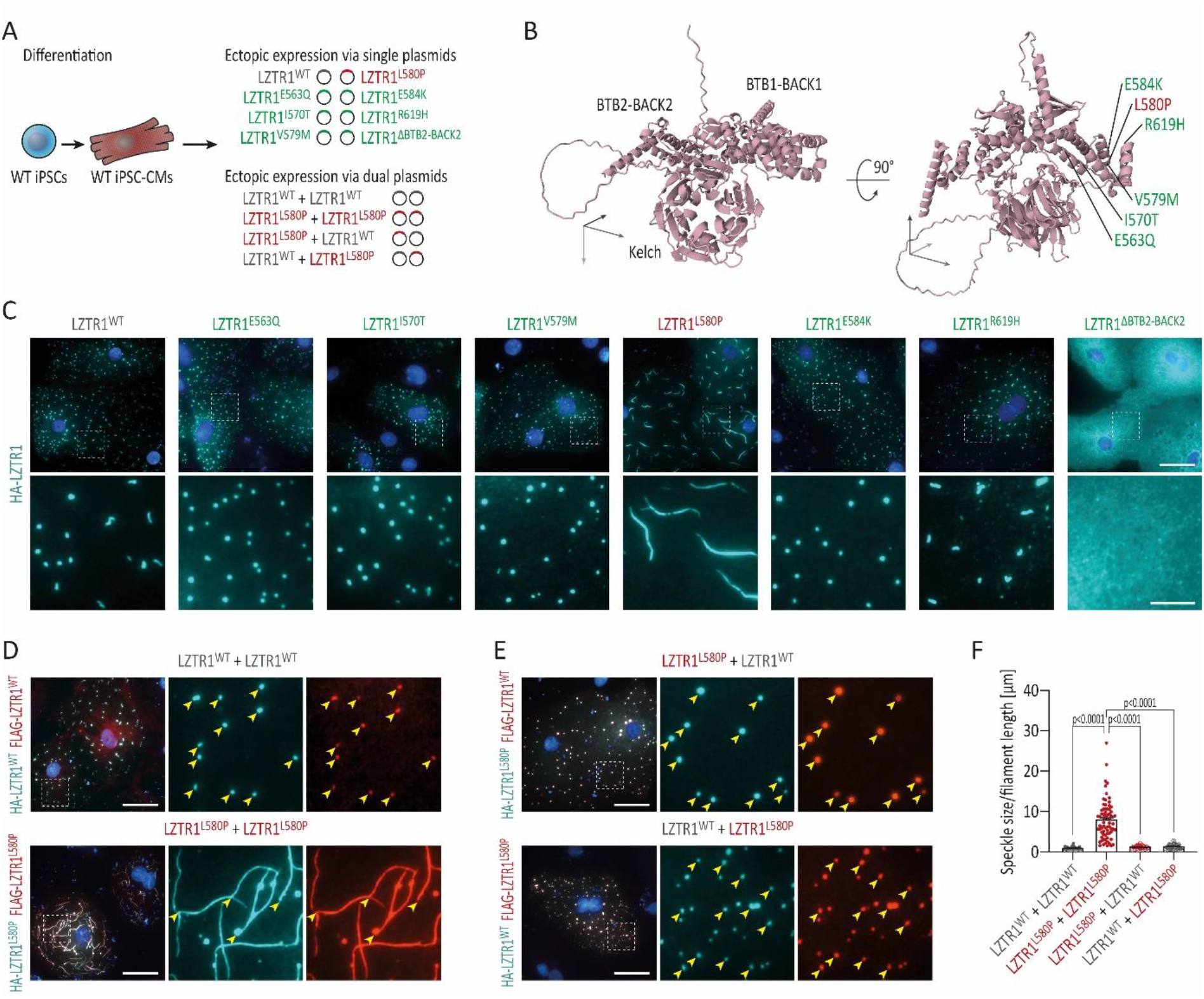
Homozygous *LZTR1^L580P^*induces polymerization of LZTR1-cullin 3 ubiquitin ligase complexes. **(A)** Depiction of the experimental design: the WT iPSC line was differentiated into ventricular iPSC-CMs, transfected at day 60 of differentiation with plasmids by lipofection for ectopic expression of LZTR1 variants and analyzed 24 h post-transfection for subcellular localization LZTR1 complexes. **(B)** AlphaFold protein structure of monomeric LZTR1 highlighting the location of selected variants within the BACK1 domain. **(C)** Representative images of iPSC-CMs after single plasmid transfection stained for HA-tagged LZTR1 revealed that LZTR1^WT^ and most other variants present a speckle-like pattern equally distributed throughout the cytoplasm, whereas missense variant LZTR1^L580P^ forms large filaments; nuclei were counter-stained with Hoechst 33342 (blue); scale bars: 20 µm in upper panel, 5 µm in lower panel. **(D-E)** Representative images of iPSC-CMs after dual plasmid transfection stained for HA-tagged and FLAG-tagged LZTR1 confirmed the filament formation of LZTR1^L580P^ (D), whereas co-expression of LZTR1^WT^ and LZTR1^L580P^ in different combinations resolved the polymer chains (E); nuclei were counter-stained with Hoechst 33342 (blue); scale bar: 20 µm. **(F)** Quantitative analysis of the mean speckle size and mean filament length per cell of HA-tagged LZTR1 in co-transfected iPSC-CMs, assessed by a customized CellProfiler pipeline, confirmed formation of *LZTR1^L580P^*-induced filaments; n=34-74 cells per condition. Data were analyzed by nonparametric Kruskal-Wallis test with Dunn correction and are presented as mean ± SEM (F).

As previously observed in other cell types (such as HeLa^8^ and HEK293^28^), LZTR1^WT^ presented as dotted pattern equally distributed throughout the cell (Figure 6C, Figure S5A in the supplement). A similar dotted appearance was observed for the variants LZTR1^E563Q^, LZTR1^I570T^, LZTR1^V579M^, LZTR1^E584K^, and LZTR1^R619H^. As expected, the truncating variant LZTR1^ΔBTB2-BACK2^ showed a mislocalized homogeneous cytoplasmic distribution. Surprisingly, missense variant LZTR1^L580P^ formed large filaments within the cytoplasm (Figure 6C, Figure S5A in the supplement). To verify this initial finding, we co-expressed two differentially tagged *LZTR1* constructs and evaluated their overlap within the cells. In accordance, LZTR1^L580P^ appeared as large protein polymers, whereas LZTR1^WT^ remained speckle-like (Figure 6D). As *LZTR1^L580P^* in heterozygous state did not induce a disease phenotype based on our clinical and our experimental evidence, we hypothesized that co-expression of LZTR1^L580P^ and LZTR1^WT^ might resolve the polymer chains. Strikingly, the LZTR1^L580P^-induced filaments dispersed when co-expressed with the WT variant, implicating that the LZTR1 complexes exclusively assembled to large protein polymers when the specific *LZTR1^L580P^* missense variant is present on both alleles (Figure 6E). To quantitatively analyze these observations, we established an automated image-based speckle/filament recognition and computation (Figure S5B in the supplement). Whereas LZTR1^WT^ displayed a mean speckle size of 0.9 µm, the mean filament length per cell in LZTR1^L580P^ amounted to 7.9 µm (Figure 6F). In line, co-expression of mutant and WT constructs, and vice versa, normalized the speckle size to 1.2 µm and 1.3 µm, respectively.

These data provide evidence that the missense variant *LZTR1^L580P^*induces a unique polymerization of LZTR1-cullin 3 ubiquitin ligase complexes, which subsequently compromises the proper function of the ubiquitination machinery.

### Homozygous LZTR1^L580P^ alters binding affinities of dimerization domains

Based on previous studies, proteins from the BTB-BACK-Kelch domain family including LZTR1 are predicted to assemble in homo-dimers.^9,28,29^ However, our current knowledge regarding the exact domains responsible for LZTR1 dimerization is limited. In order to identify a plausible explanation for the *LZTR1^L580P^*-induced polymerization, we utilized ColabFold – an AlphaFold-based platform for the prediction of protein structures and homo-and heteromer complexes.^30^ We used a homo-trimer configuration of the experimentally employed *LZTR1* variants (all within the BACK1 domain) and AlphaFold-multimer predicted five high-quality models each with an average predicted local distance difference test (a per-residue confidence metric) between 64.1 and 76.0. For all variants, we inspected the interaction between the chains through the predicted alignment error (PAE) generated by AlphaFold-multimer (Figure S6 in the supplement). Here, a low PAE indicates that the relative position and orientation of the positions x and y was correctly predicted – a measure indicating if interfacing residues were correctly predicted across chains. Based on these predictions, we compared the top-ranked models of each variant according to the predicted template modeling score, which corresponded to overall topological accuracy (Figure 7A). The top-ranked model for LZTR1^WT^ showed interaction as a homo-dimer via the BACK2-BACK2 domain, whereas the third LZTR1 protein remained monomeric. We also observed the identical dimerization via the BACK2 domains for all other variants, except for LZTR1^L580P^ (Figure S6 in the supplement). In contrast, the top-ranked model for LZTR1^L580P^ predicted an interaction between all three chains, on the one hand via the BACK2-BACK2 domain and on the other hand via the BACK1-BACK1 domain (Figure 7A). In addition, we used AlphaFold-multimer to predict the interaction of LZTR1^L580P^ with the substrate MRAS and the ubiquitin ligase cullin 3 (Figure 7B). Within the multiprotein complex, MRAS was predicted to bind to the Kelch domain, whereas cullin 3 was predicted to interact with the BTB1-BTB2 domain of LZTR1.

**Figure 7:**
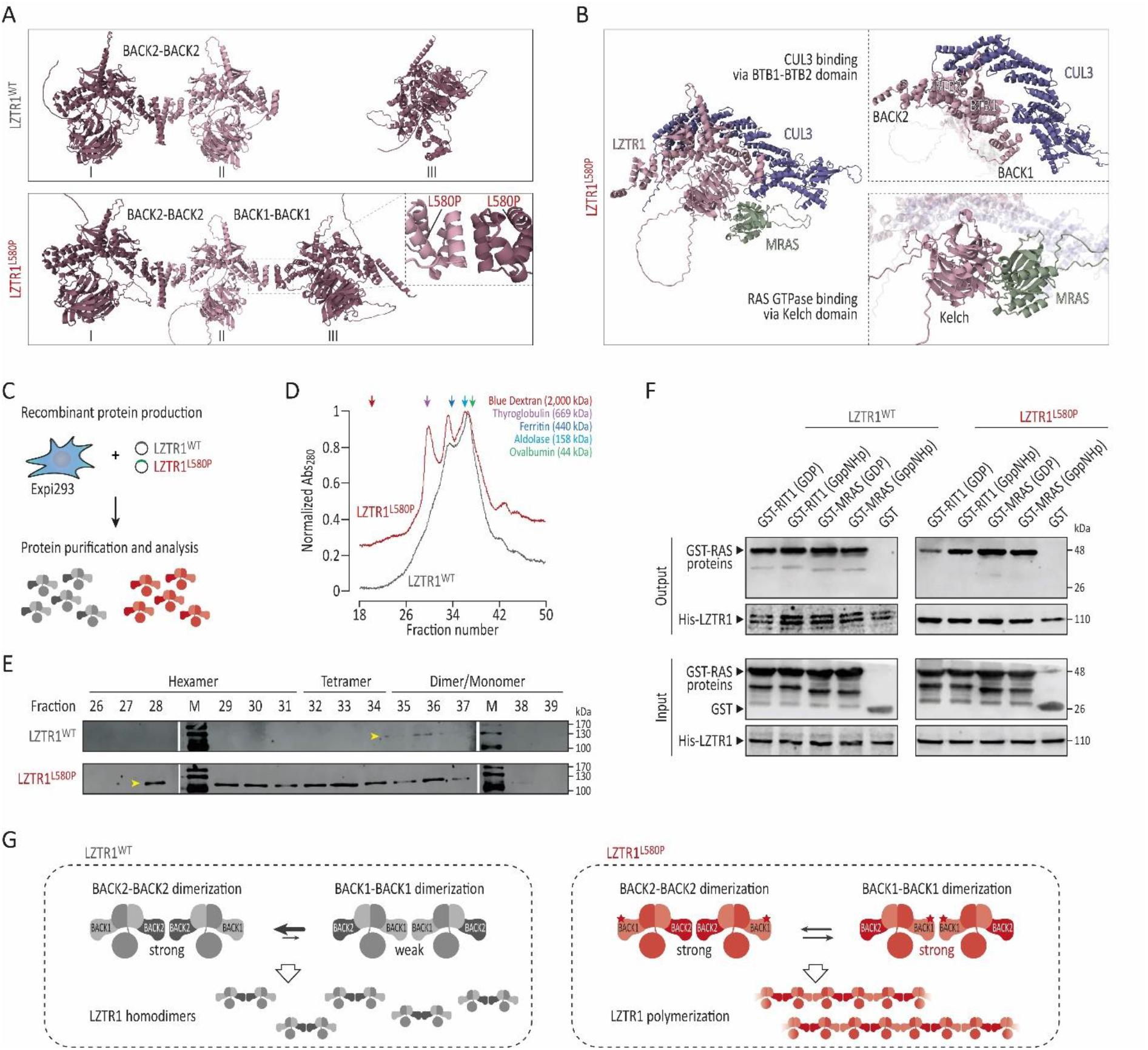
Homozygous *LZTR1^L580P^*alters binding affinities of dimerization domains. **(A)** Computational modeling of the top-ranked LZTR1 homo-trimer interactions of selected variants within the BACK1 domain, assessed by the predicted alignment error generated by ColabFold, predicted a dimer plus monomer configuration via BACK2-BACK2 dimerization for LZTR1^WT^ and the other variants, whereas the top-ranked model for LZTR1^L580P^ was predicted to form linear trimers via BACK2-BACK2 and BACK1-BACK1 dimerization. **(B)** Computational modeling of the interaction between LZTR1^L580P^ and its binding partners predicted binding to cullin 3 (CUL3) via the BTB1-BTB2 domain and to MRAS via the Kelch domain. **(C)** Production of LZTR1^WT^ and LZTR1^L580P^ recombinant proteins from Expi-293F cells for characterization of molecular masses of proteins and protein complexes. **(D)** Analytical size exclusion chromatography of soluble recombinant LZTR1 proteins revealed a higher order oligomerization profile for LZTR1^L580P^ compared to the less complex elution profile of LZTR1^WT^. **(E)** Immunoblotting of the fractions showed elution of LZTR1^L580P^ as hexamer, tetramer, and dimer/monomer, whereas LZTR1^WT^ eluted predominantly as dimer/monomer. **(F)** Pull-down assay analysis showed comparable binding affinities of LZTR1^WT^ and LZTR1^L580P^ with MRAS and RIT1 proteins in both inactive (GDP-bound) and active (GppNHp-bound) states. **(G)** Hypothetical model for LZTR1 complex formation: whereas LZTR1^WT^ assembles in homo-dimers via the BACK2-BACK2 dimerization domain, LZTR1^L580P^ might alter the binding affinity of the BACK1 domain, causing formation of linear LZTR1 polymer chains via dimerization of both BACK2 and BACK1 domains.

To experimentally confirm the formation of LZTR1^L580P^ polymers, we produced soluble recombinant proteins of LZTR1^WT^ and LZTR1^L580P^ and analyzed the purified samples by analytical size exclusion chromatography, allowing to characterize the molecular masses of proteins and protein complexes (Figure 7C). A higher order oligomerization profile was observed for LZTR1^L580P^, whereas LZTR1^WT^ exhibited a less complex elution profile (Figure 7D). Immunoblotting of the fractions showed that LZTR1^L580P^ eluted as a hexamer with a molecular weight of approximately 700 kDa, as a tetramer corresponding to 450-550 kDa, and as a dimer/monomer with a molecular weight of 100-200 kDa (Figure 7E). In contrast, LZTR1^WT^ was characterized by a single peak, indicative of its predominantly dimeric/monomeric state. In addition, we examined the interaction of LZTR1^WT^ and LZTR1^L580P^ proteins with RIT1 and MRAS in their inactive (GDP-bound) and active (GppNHp-bound; GppNHp is a non-hydrolyzable GTP-analog) states. Both LZTR1^WT^ and mutant LZTR1^L580P^ were capable of binding their substrates in both nucleotide-bound states (Figure 7F).

Collectively, the *in silico* predictions and molecular analyses suggest that the missense variant *LZTR1^L580P^* alters the binding affinities of the BACK1 domain enabling formation of linear LZTR1 polymer chains via both dimerization domains, thereby providing a rationale for the molecular and cellular impairments in NS (Figure 7G).

### Homozygous LZTR1^L580P^ preserves residual function of ubiquitin ligase complex

To investigate how severely the degradation of RAS GTPases is affected by the missense variant LZTR1^L580P^ and the polymerization of LZTR1 complexes (especially compared to the complete loss of LZTR1), we treated the patient-specific iPSC-CMs, the CRISPR-corrected cells, the LZTR1^KO^ cells, and the WT controls with the cullin RING ligase inhibitor pevonedistat (which blocks the ubiquitin-mediated degradation via the proteasome and the autophagosome) or the proteasome inhibitor MG-132 and analyzed MRAS and RIT1 protein levels three days after treatment (Figure 8A,B). As expected, inhibition of cullin-mediated ubiquitination by pevonedistat increased MRAS and RIT1 protein levels in WT and CRISPR-corrected iPSC-CMs (Figure 8C-E). Treatment in patient-specific LZTR1^L580P^ cultures further increased the RAS GTPase levels, indicating residual function of the LZTR1^L580P^-cullin 3 ubiquitin ligase complex. Interestingly, while MRAS accumulation in LZTR1^KO^ cultures could not be further increased by cullin inhibition, RIT1 protein levels were significantly higher after treatment in *LZTR1*-deficient cells. This suggests that MRAS is exclusively targeted for degradation by the LZTR1-cullin 3 ubiquitin ligase complex, whereas RIT1 can be additionally degraded in an *LZTR1*-independent manner. Furthermore, inhibition of the ubiquitin-proteasome system resulted in increased RIT1 levels, suggesting that RIT1 is predominantly degraded by the proteasomal pathway (Figure 8F-H). In contrast, MRAS levels were not affected after treatment across all iPSC lines, indicating exclusive autophagosomal degradation of MRAS.

**Figure 8:**
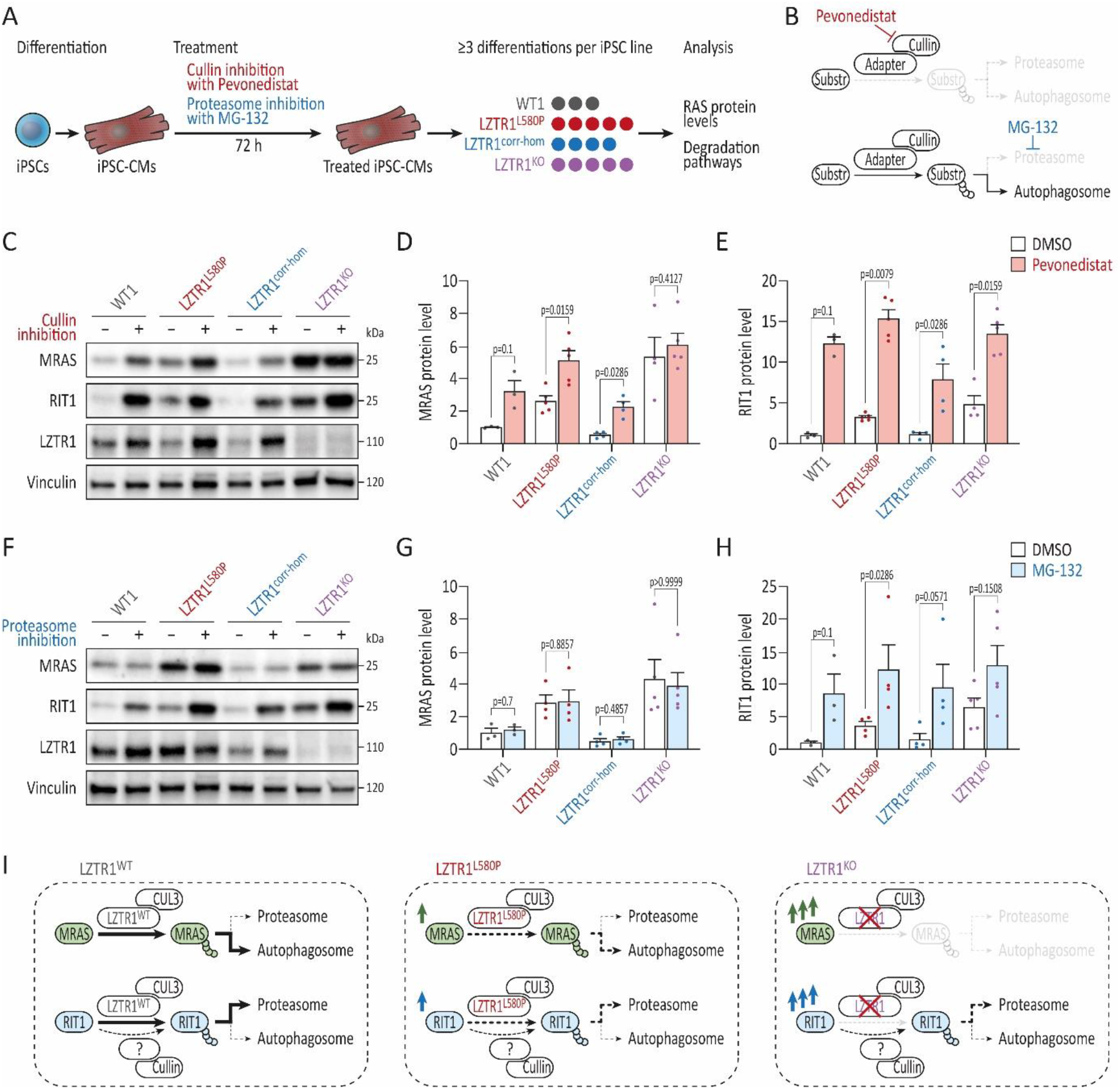
Homozygous *LZTR1^L580P^*preserves residual function of ubiquitin ligase complex. **(A)** Depiction of the experimental design: the WT, the patient-specific, the homozygous CRISPR-corrected, and LZTR1^KO^ iPSC lines were differentiated into ventricular iPSC-CMs and treated with pevonedistat and MG-132 for 3 days to analyze the ubiquitin-mediated degradation of RAS GTPases; n=3-5 individual differentiations/treatments per iPSC line. **(B)** Mode of action of pevonedistat and MG-132 on degradation pathways: pevonedistat is a selective NEDD8-activating enzyme inhibitor, preventing neddylation of cullin RING ligases and blocking ubiquitin-mediated degradation via the proteasome and the autophagosome, whereas MG-132 is a selective inhibitor specifically blocking the proteolytic activity of the 26S proteasome. **(C)** Representative blots showing MRAS, RIT1, and LZTR1 levels in WT, patient’s, CRISPR-corrected, and LZTR1^KO^ iPSC-CMs upon pevonedistat treatment for 3 days, assessed by Western blot; Vinculin served as loading control. **(D-E)** Quantitative analysis of Western blots for MRAS (D) and RIT1 (E) upon pevonedistat treatment; data were normalized to total protein and to the DMSO-treated WT samples on each membrane. **(F)** Representative blots showing MRAS, RIT1 and LZTR1 levels in WT, patient’s, CRISPR-corrected, and LZTR1^KO^ iPSC-CMs upon MG-132 treatment for 3 days, assessed by Western blot; Vinculin served as loading control. **(G-H)** Quantitative analysis of Western blots for MRAS (G) and RIT1 (H) upon MG-132 treatment; data were normalized to total protein and to the DMSO-treated WT samples on each membrane. Data were analyzed by nonparametric Kruskal-Wallis test with Dunn correction and are presented as mean ± SEM (D, E, G, H). **(I)** Proposed model for LZTR1-mediated degradation of MRAS and RIT1 for LZTR1^WT^, LZTR1^L580P^, and LZTR1^KO^: MRAS is exclusively targeted by the LZTR1^WT^-cullin 3 ubiquitin ligase complex for degradation via autophagy, whereas RIT1 is additionally ubiquitinated by other cullin ubiquitin ligases and degraded predominantly by the proteasome; b) the LZTR1^L580P^ decreases degradation of MRAS and RIT1; c) loss of LZTR1 completely prevents MRAS degradation, while RIT1 degradation remains to some extent in an LZTR1-independent manner.

These data confirm that the missense variant *LZTR1^L580P^*preserves some residual function of the LZTR1-cullin 3 ubiquitin ligase complex compared to the complete loss of *LZTR1*. Furthermore, the results demonstrate that degradation of cardiomyocyte-specific MRAS is exclusively mediated by LZTR1 via the autophagosome, whereas proteasomal degradation of RIT1 is mediated by both *LZTR1*-dependent and *LZTR1*-independent pathways.

## Discussion

Both autosomal dominant and autosomal recessive forms of *LZTR1*-associated NS have been described presenting with a broad clinical spectrum and various phenotypic expression of symptoms. However, the mechanistic consequences of numerous of these mutations, mostly classified as variants of uncertain significance, are still under debate. In previous studies, we and others elucidated the role of LZTR1 as a critical negative regulator of the RAS-MAPK pathway by controlling the pool of RAS GTPases.^8–10,28,31^ By using patient-derived iPSC-CMs from NS patients with biallelic truncating *LZTR1* variants, we could show that *LZTR1* deficiency results in accumulation of RAS levels, signaling hyperactivity, and cardiomyocyte hypertrophy.^10^ Further, by genetically correcting one of the two affected alleles, we could show that one functional *LZTR1* allele is sufficient to maintain normal RAS-MAPK activity in cardiac cells. In contrast to the truncating variants, dominant *LZTR1* missense variants generally cluster in the Kelch motif. Based on heterologous expression systems, these dominant variants are considered to perturb recognition or binding of RAS substrates to the LZTR1 ubiquitination complex.^8,9,11,31^ Much less is known about the functional relevance of recessive *LZTR1* missense variants, which are distributed over the entire protein. Detailed insights in the underlying molecular and functional mechanisms of selective variants causing the severe cardiac phenotype in NS enable to gain insights into specific structure-function relations of LZTR1 and are crucial to facilitate the development of patient-specific therapies.

In this study, we diagnosed a patient with NS, who presented typical clinical features of NS including an early-onset HCM and confirmed this diagnosis on genetic level by the identification of the homozygous, causative variant c.1739T>C/p.L580P in *LZTR1* by whole exome sequencing. The variant is novel and was not described before in patients with NS and we classified *LZTR1^L580P^*as likely causative based on its absence in gnomAD and our computational prediction. Besides the *LZTR1* variant, no additional variants were detected in other NS-associated genes or novel RAS-associated candidate genes. By combining *in vitro* disease modeling using patient-specific and CRISPR/Cas9-corrected iPSC-CMs, with molecular and cellular phenotyping, as well as *in silico* structural modeling, we uncovered a unique *LZTR1^L580P^*-specific disease mechanism provoking the cardiac pathology of NS. In detail, we found that a) *LZTR1^L580P^* is predicted to alter the binding affinity of the BACK1 dimerization domain facilitating the formation of linear LZTR1 protein chains; b) homozygous *LZTR1^L580P^*fosters the assembly of large polymers of LZTR1-cullin 3 ubiquitin ligase complexes; c) pathological polymerization results in LZTR1 complex dysfunction, disturbed ubiquitination, accumulation of RAS GTPases, and RAS-MAPK signaling hyperactivity; and d) increased signaling activity induces global changes of the proteomic landscape ultimately causing cellular hypertrophy. Importantly, correction of one allele – in line with co-expression of WT and mutant *LZTR1* transcripts – is sufficient to normalize the cardiac disease phenotype both on molecular and cellular level.

Based on recent publications, there is a broad consensus about the role of LZTR1 as an adaptor protein for the cullin 3 ubiquitin ligase complex targeting RAS proteins for ubiquitination and subsequent protein degradation.^8–11,28,31^ In line with observations in other NS-associated genes and mutations, LZTR1 dysfunction and concomitant accumulation of RAS GTPases results in hyperactivation of RAS-MAPK signaling. We confirmed robustly elevated RAS levels in patient-specific cells harboring the homozygous *LZTR1^L580P^* missense variant. However, the accumulation of RAS GTPases and ERK hyperactivity was substantially higher in LZTR1^KO^ cells, supporting a partial residual function of LZTR1^L580P^ ubiquitin ligase complexes. Furthermore, it remains controversial, whether LZTR1 is able to recognize all members of the RAS GTPase family for degradation or whether there is a selective affinity towards particular RAS members. By using heterologous expression systems, LZTR1 interaction with the main highly conserved RAS proteins HRAS, KRAS and NRAS was observed.^8,28,31^ On the contrary, Castel and colleagues observed a selective binding of LZTR1 with RIT1 and MRAS, but not with HRAS, KRAS or NRAS.^9^ Moreover, in homozygous *LZTR1* knockout mice elevated RIT1 protein levels were detected across different organs including brain, liver, and heart, whereas HRAS, KRAS, and NRAS levels (recognized by pan-RAS) remained unchanged.^32^ By using global proteomics, we now provide further evidence that LZTR1 dysfunction in cardiomyocytes in particular causes severe accumulation of MRAS and RIT1 and, to a lower extent, upregulation of the other RAS GTPases HRAS, KRAS and NRAS, although all RAS proteins are robustly expressed in this cell type. We conclude that based on gene expression data and overall protein levels, MRAS seems to be the most prominent RAS candidate in cardiomyocytes driving the signaling hyperactivity in these cells. In addition, our inhibition experiments demonstrate that MRAS degradation is exclusively mediated by LZTR1 via the autophagosome, whereas RIT1 degradation is mediated by both LZTR1-dependent and LZTR1-independent pathways. These observations suggest that at endogenous expression levels, LZTR1 possesses a certain selectivity for MRAS and RIT1, and a lower affinity for the RAS GTPases HRAS, KRAS and NRAS. However, we cannot exclude the possibility of cell type-specific differences in LZTR1-RAS binding affinities.

Besides accumulation of RAS members, HSPA2 was strongly upregulated in *LZTR1*-deficient iPSC-CMs both on transcriptional as well as on protein level, suggesting that HSPA2 is not a direct substrate of LZTR1. In line, severely increased HSPA2 levels had been observed by us in NS iPSC-CMs with *LZTR1*-truncating variants,^10^ in a *RAF1*-related NS iPSC-CM model,^15^ as well as in iPSC-CMs from Fabry disease patients, a lysosomal storage disorder associated with cardiac involvement such as HCM and arrhythmias.^33^ Moreover, a significant cardiomyocyte-specific elevation of HSPA2 was also observed in HCM tissue from patients.^34,35^ A heat shock protein 70-based therapy has been shown to reverse lysosomal pathology,^36^ whereas deletion of these gene members was assumed to induce cardiac dysfunction and development of cardiac hypertrophy.^37^ As a member of the large group of chaperones, HSPA2 is known to have a dual function in cells: to mediate disaggregation and refolding of misfolded proteins as well as to assist in protein degradation via the ubiquitin-proteasome system or the lysosome-mediated autophagy.^38,39^ This suggests that HSPA2 upregulation may be a cardio-protective adaptive response of the hypertrophic cardiomyocytes to cope with *LZTR1^L580P^*-induced RAS accumulation (and possibly LZTR1 polymerization), by regulating the quality control mechanisms for protein degradation.

Major hallmarks of pathological cardiac hypertrophy include impaired cardiac function, changes in extracellular matrix composition, and fibrosis, as well as metabolic reprogramming and mitochondrial dysfunction.^40^ In accordance, the proteomic disease signature of patient-derived iPSC-CM cultures revealed impairments in muscle contraction, extracellular matrix organization, and metabolism, all crucial for proper cardiomyocyte function. Furthermore, *LZTR1^L580P^*-derived iPSC-CMs recapitulated the patient’s hypertrophic phenotype reflected by cellular enlargement. Strikingly, both the molecular profile as well as cellular hypertrophy were resolved upon CRISPR-correction of the missense variant. Interestingly, no myofibrillar disarray was observed in our cell model. However, the presence of myofibril disarray in NS remains controversial: whereas structural defects were described in *RAF1*-associated iPSC models,^15,41^ we and others did not observe any impact on sarcomere structures or myofibril organization in *LZTR1*-related, *PTPN11*-related, and *BRAF*-related iPSC-CMs,^10,14,16^ implying potential genotype-dependent differences in the manifestation of myofibril disassembly in NS. Missense variants in *LZTR1* located within the Kelch domain are predicted to affect substrate recognition, whereas missense variants in the BTB-BACK domain are assumed to impair either binding of cullin 3, proper homo-dimerization, or correct subcellular localization. Several studies could provide proof that dominantly acting Kelch domain variants perturb recognition of RAS substrates, but do not affect LZTR1 complex stability or subcellular localization.^8,9,11,31,42^ Vice versa, BTB-BACK missense variants showed no influence on RIT1 binding.^9,42^ However, variants located in the BTB1 or the BTB2 domain, such as *LZTR1^V456G^*, *LZTR1^R466Q^*, *LZTR1^P520L^*, and *LZTR1^R688C^*, caused a subcellular mislocalization from defined speckles to a diffuse cytoplasmic distribution, similar to the findings obtained from truncating *LZTR1* variants.^8,31^ In addition to these distinct pathological consequences from different variants analyzed so far, we now provide evidence for an alternative disease mechanism unique to BACK1 domain-located *LZTR1^L580P^*: ectopic expression of *LZTR1^L580P^* in iPSC-CMs caused a pathological polymerization of LZTR1 ubiquitination complexes. This phenomenon was verified by *in silico* prediction and chromatography with purified recombinant LZTR1 proteins. In contrast, the binding probabilities of *LZTR1^L580P^* to substrates and interaction partners were not significantly affected by the mutation. This remarkable phenotype was not observed for any other variant within the BACK1 domain. Notably, ectopic co-expression of *LZTR1^L580P^*and *LZTR1^WT^* alleviated the polymerization, indicating that the assembly of LZTR1 polymer chains exclusively occurs if the mutated proteins are present in homozygous state. Strikingly, an oligomerization of another BTB-BACK family member had been reported previously: Marzahn and colleagues revealed that dimers from the cullin 3 ubiquitin ligase substrate adaptor SPOP (harboring only one BTB-BACK domain) self-associate into linear higher-order oligomers via BACK domain dimerization.^43^ These SPOP oligomers assembled in membrane-less cellular bodies, visualized as nuclear speckles, and it was proposed that the speckles might be important hotspots of ubiquitination. Based on these findings and our data, we propose that LZTR1 complexes concentrate in cellular speckles (either as dimers or as oligomers) to form subcellular clusters for efficient ubiquitination and degradation of RAS proteins. However, *LZTR1^L580P^*-induced polymerization of these complexes compromises regular function, leading to accumulation of substrates. The CRISPR-based correction was able to rescue the polymerization phenotype and may be a sustainable treatment option in the future. Alternatively, it may be possible to identify compounds that specifically prevent the interaction of LZTR1 complexes via BACK1-BACK1 dimerization.

Our knowledge about the particular domains responsible for LZTR1 homo-dimerization is still incomplete. Whereas Castel and colleagues proposed that the BTB1 and the BACK1 domain are required for dimerization,^9^ Steklov et al. observed impaired assembly in a BACK2 domain mutated *LZTR1* variant.^31^ Based on *in silico* modeling, we now propose that LZTR1 can dimerize either via the BACK2-BACK2 domains or via the BACK1-BACK1 domains. Although in LZTR1^WT^ the BACK2-BACK2 dimerization might be primarily utilized, changes in binding affinities of the BACK1 domain as a consequence of *LZTR1^L580P^*facilitated tandem self-association of dimers to linear multimers. Strikingly, the *in silico* models for complex assembly of the different variants were consistent with the experimental data. However, this analysis must be taken with caution as the PAE signal across chains is overall weak and AlphaFold-multimer was not trained with single point variants in mind. Of note, dimer/monomer as well as trimer interactions (in diverse combinations, such as via BACK2-BACK2 and BACK1-BACK1 or via BACK2-BACK2-BACK2) had also been predicted for the other BACK1 variants as well as for WT in the lower-ranked models (Figure S6 in the supplement). As a next step, it would be interesting to see, if the trend stays consistent for complexes with more chains. However, due to technical prerequisites, we were currently not able to predict more than three chains.

So far, the relevance of certain *LZTR1* missense variants on the molecular and cellular processes had been investigated in heterologous expression systems, failing to faithfully represent human cardiac physiology. Our study demonstrates the potential of patient-specific iPSCs to model human diseases and to uncover variant-specific pathomechanisms, which might facilitate the development for early and more precise therapies. Despite the great advantages of this model system over other cellular models, iPSC-CMs possess certain limitations. As summarized by several reports, iPSC-CMs are considered to be developmentally immature characterized by molecular and functional properties similar to fetal CMs.^25,44,45^ Although we complemented our study by utilizing three-dimensional EHMs, these *in vitro* models are currently not able to entirely resemble the disease phenotype at organ level. Nevertheless, our investigations at single cell and tissue level have proven to be a valuable platform for uncovering disease-relevant signaling pathways, identifying novel therapeutic targets and studying the disease progression during cardiogenesis.

Taken together, this study uncovered a novel mechanism causing recessive NS, which is initiated by *LZTR1^L580P^*-driven polymerization of LZTR1 ubiquitination complexes, provoking molecular and cellular impairments associated with cardiac hypertrophy. Moreover, CRISPR-correction of the missense variant on one allele was sufficient to rescue the phenotype, thereby providing proof-of-concept for a sustainable therapeutic approach.

## Materials and methods

### Ethical approval

The study was approved by the Ethics Committee of the University Medical Center Göttingen (approval number: 10/9/15) and carried out in accordance with the approved guidelines. Written informed consent was obtained from all participants or their legal representatives prior to the participation in the study.

### Whole exome sequencing

Whole exome sequencing on genomic DNA of the patient was performed using the SureSelect Human All Exon V6 kit (Agilent) on an Illumina HiSeq 4000 sequencer. The “Varbank 2.0” pipeline of the Cologne Center for Genomics (CCG) was used to analyze and interpret the exome data, as previously described.^10^ Co-segregation analysis was performed in the family. Computational predictions for the pathogenicity of the variant were performed using MutationTaster (https://www.mutationtaster.org/), SIFT (https://sift.bii.a-star.edu.sg/), and PolyPhen-2 (http://genetics.bwh.harvard.edu/pph2/).

### Generation and culture of human iPSCs

Human iPSC lines from two healthy donors, from one NS patient with biallelic truncating variants in *LZTR1* (NM_006767.4: c.27dupG/p.Q10Afs*24, c.1943-256C>T/p.T648fs*36), from one NS patient with a pathological missense variant in *LZTR1* (NM_006767.4: c.1739T>C/p.L580P; ClinVar: RCV000677201.1), as well as heterozygous and homozygous CRISPR/Cas9-corrected iPSC lines were used in this study. Wild type iPSC lines UMGi014-C clone 14 (isWT1.14, here abbreviated as WT1) and UMGi130-A clone 8 (isWT11.8, here abbreviated as WT11) were generated from dermal fibroblasts and peripheral blood mononuclear cells from two male donors, respectively, using the integration-free Sendai virus and described previously.^46,47^ Patient-specific iPSC line UMGi030-A clone 14 (isHOCMx1.14, here abbreviated as LZTR1^KO^) was generated from patient’s dermal fibroblasts using the integration-free Sendai virus and described previously.^10^ Patient-specific iPSC line UMGi137-A clone 2 (isNoonSf1.2, here abbreviated as LZTR1^L580P^) was generated from patient’s dermal fibroblasts using the integration-free Sendai virus according manufacturer’s instructions with modifications, as previously described.^10^ Genetic correction of the pathological gene variant in the patient-derived iPSC line UMGi137-A clone 2 was performed using ribonucleoprotein-based CRISPR/Cas9 using crRNA/tracrRNA and Hifi SpCas9 (IDT DNA technologies) by targeting exon 15 of the *LZTR1* gene, as previously described.^10^ The guide RNA target sequence was (PAM in bold): 5’-GCGGCACTCTCGCACACAAC **CGG**-3’. For homology-directed repair, a single-stranded oligonucleotide with 45-bp homology arms was used. After automated clonal singularization using the single cell dispenser CellenOne (Cellenion/Scienion) in StemFlex medium (Thermo Fisher Scientific), successful genome editing was identified by Sanger sequencing and the CRISPR-corrected isogenic iPSC lines UMGi137-A-1 clone D8 (isNoonSf1-corr.D8, here abbreviated as L580P^corr-het^) and UMGi137-A-1 clone D1 (isNoonSf1-corr.D1, here abbreviated as L580P^corr-hom^) were established. Newly generated iPSC lines were maintained on Matrigel-coated (growth factor reduced, BD Biosciences) plates, passaged every 4-6 days with Versene solution (Thermo Fisher Scientific) and cultured in StemMACS iPS-Brew XF medium (Miltenyi Biotec) supplemented with 2 µM Thiazovivin (Merck Millipore) on the first day after passaging with daily medium change for at least ten passages before being used for molecular karyotyping, pluripotency characterization, and differentiation experiments. Pluripotency analysis via immunocytochemistry and flow cytometry was performed, as previously described.^10^ For molecular karyotyping, genomic DNA of iPSC clones was sent for genome-wide analysis via Illumina BeadArray (Life&Brain, Germany). Digital karyotypes were analyzed in GenomeStudio v2.0 software (Illumina). For off-target screening, the top five predicted off-target regions for the respective guide RNA ranked by the CFD off-target score using CRISPOR^48^ were analyzed by Sanger sequencing. Human iPSCs and iPSC-derivatives were cultured in feeder-free and serum-free culture conditions in a humidified incubator at 37°C and 5% CO_2_. All antibodies used for immunofluorescence and flow cytometry are listed in Table S2 in the supplement.

### Cardiomyocyte differentiation of iPSCs and generation of engineered heart muscle

Human iPSC lines were differentiated into ventricular iPSC-CMs via WNT signaling modulation and subsequent metabolic selection, as previously described,^19^ and cultivated in feeder-free and serum-free culture conditions until day 60 post-differentiation before being used for molecular and cellular experiments. Defined, serum-free EHMs were generated from iPSC-CMs around day 30 of differentiation and human foreskin fibroblasts (ATCC) at a 70:30 ratio according to previously published protocols.^26^ Optical analysis of contractility and rhythm of spontaneously beating EHMs in a 48 well plate (myrPlate TM5, myriamed GmbH) was performed between day 34 and day 42 of culture using a custom-built setup with a high-speed camera by recording the movement of the two UV light-absorbing flexible poles. Contractility parameters of EHM recordings of at least 1 min recording time were analyzed via a custom-build script in MatLab (MathWorks). For each iPSC line, three individual differentiations were used for EHM casting.

### Biosensor-based analysis of ERK signaling dynamics in iPSC-CMs

In brief, the ERK kinase translocation reporter (ERK-KTR) biosensor consists of an ERK-specific docking site, a nuclear localization signal (NLS), a nuclear export signal (NES), and mClover. Endogenous, phosphorylated ERK binds to the biosensor and phosphorylates its NLS and NES resulting in a nucleus-cytoplasm shuttling according to ERK activity.^20^ ERK-KTR biosensor encoding lentiviral particles were produced in HEK293T cells transfected with transfer, envelope, and packaging plasmids using Lipofectamine 3000 (Thermo Fisher Scientific) according to manufacturer’s instructions. pLentiPGK Puro DEST ERKKTRClover was a gift from Markus Covert (RRID:Addgene_90227), pMD2.G was a gift from Didier Trono (RRID:Addgene_12259), and psPAX2 was a gift from Didier Trono (RRID:Addgene_12260). Virus was harvested from day 2 to day 5 post-transfection by medium collection and centrifugation at 500×g at 4°C for 5 min. The harvested virus was filtered using a 0.45 µm filter and a syringe. For transduction, 15,000 iPSC-CMs were seeded per well of a 96-well plate and lentiviral transduction was performed 7 days after cell digestion. Lentivirus was diluted in culture medium supplemented with 100 U/ml penicillin, 100 µg/ml streptomycin (Thermo Fisher Scientific), and 10 µg/ml Polybrene Transfection Reagent (Merck). After 24 h of incubation, medium was replaced with cardio culture medium and cells were maintained for additional 7 days. For live-cell imaging, biosensor-transduced iPSC-CM cultures at day 60 of differentiation were treated with 100 nM MEK inhibitor trametinib (Selleck Chemicals), 100 nM JNK inhibitor JNK-IN-8 (Hycultec), or 1:1,000 DMSO (Sigma-Aldrich) for 60 min, before stimulation with 10% fetal bovine serum for another 60 min. Cells were imaged every 10 min for a total time of 120 min. Live cell imaging experiments were acquired using the CQ1 confocal image cytometer (Yokogawa Electric Corporation) and CellPathfinder software (Yokogawa Electric Corporation) under environmental control (37°C, 5% CO_2_). Exported images were processed using the StarDist method for nucleus segmentation.^49^ The StarDist network was retrained on 60 images from our dataset with annotations manually produced with napari.^50^ For a new image, the nuclei were then segmented with StarDist, and a ring element around each nucleus was computed to approximate the cytosol. The mean fluorescence intensity of both compartments was measured for each cell individually.

### Proteomics and Western blot analysis of iPSC-CMs

For proteomic analysis, iPSC-CMs were pelleted at day 60 of differentiation by scratching in RIPA buffer (Thermo Fisher Scientific) containing phosphatase and protease inhibitor (Thermo Fisher Scientific) and snap-frozen in liquid nitrogen. Cell pellets were reconstituted in 8 M urea/ 2 M thiourea solution (Sigma-Aldrich) and lysed by five freeze-thaw cycles at 30°C and 1.600 rpm. Protein containing supernatant was collected by centrifugation. Nucleic acid was degraded enzymatically with 0.125 U/µg benzonase (Sigma-Aldrich), and protein concentration was determined by Bradford assay (Bio-Rad). Five µg protein was processed for LC-MS/MS analysis, as previously described.^51^ Briefly, protein was reduced (2.5 mM dithiothreitol, Sigma-Aldrich; 30 min at 37°C) and alkylated (10 mM iodacetamide, Sigma-Aldrich; 15 min at 37°C) before proteolytic digestion with LysC (enzyme to protein ratio 1:100, Promega) for 3 h and with trypsin (1:25, Promega) for 16 h both at 37°C. The reaction was stopped with 1% acetic acid (Sigma-Aldrich), and the peptide mixtures were desalted on C-18 reverse phase material (ZipTip μ-C18, Millipore). Eluted peptides were concentrated by evaporation under vacuum and subsequently resolved in 0.1% acetic acid / 2% acetonitrile containing HRM/iRT peptides (Biognosys) according to manufacturer’s recommendation. LC-MS/MS analysis was performed in data-independent acquisition (DIA) mode using an Ultimate 3000 UPLC system coupled to an Exploris 480 mass spectrometer (Thermo Scientific). Peptides were separated on a 25 cm Accucore column (75 µm inner diameter, 2.6 µm, 150 A, C18) at a flow rate of 300 nl/min in a linear gradient for 60 min. Spectronaut software (Biognosys) was used for the analysis of mass spectrometric raw data. For peptide and protein identification, the Direct DIA approach based on UniProt database limited to human entries was applied. Carbamidomethylation at cysteine was set as static modification, oxidation at methionine and protein N-terminal acetylation were defined as variable modifications, and up to two missed cleavages were allowed. Ion values were parsed when at least 20% of the samples contained high quality measured values. Peptides were assigned to protein groups and protein inference was resolved by the automatic workflow implemented in Spectronaut. Statistical data analysis was conducted using an in-house developed R tool and based on median-normalized ion peak area intensities. Methionine oxidized peptides were removed before quantification. Differential abundant proteins (p-value ≤ 0.05) were identified by the algorithm ROPECA^52^ and application of the reproducibility-optimized peptide change averaging approach^53^ applied on peptide level. Only proteins quantified by at least two peptides were considered for further analysis. Reactome pathway enrichment analysis was performed using the ClueGo plugin in Cytoscape.^54^ For each iPSC line, at least three individual differentiations were analyzed. For Western blot analysis, protein containing supernatant was collected by centrifugation. Protein concentration was determined by BCA assay (Thermo Fisher Scientific). Samples were denatured at 95°C for 5 min. 15 µg protein were loaded onto a 4-15% Mini-PROTEAN TGX Stain-Free precast gel (Bio-Rad). The protein was separated by sodium dodecyl sulfate-polyacrylamide gel electrophoresis (SDS-PAGE) by applying 200 V for 30 min. Post-running, TGX gels were activated via UV light application using the Trans-Blot Turbo transfer system (Bio-Rad). While blotting, proteins were transferred to a nitrocellulose membrane (25 V constant, 1.3 A for 7 min). Total protein amount was detected via the ChemiDoc XRS+ (Bio-Rad) system and used for protein normalization. After 1 h in blocking solution (5% milk in TBS-T, Sigma-Aldrich), membranes were incubated in primary antibody solution (1% milk in TBS-T) overnight. Membrane was washed trice with TBS-T before applying the secondary antibody (1:10,000 in 1% milk in TBS-T) at RT for 1 h. After washing, signals were detected upon application of SuperSignal West Femto Maximum Sensitivity Substrat (Thermo Fisher Scientific). Image acquisition was performed with the ChemiDoc XRS+ (Bio-Rad) at the high-resolution mode. For protein quantification, ImageLab (Bio-Rad) was used and protein levels were first normalized to total protein and second to the corresponding WT samples on each blot. For ERK signaling analysis, iPSC-CMs at day 60 of differentiation were treated with 10 nM trametinib (Selleck Chemicals) for 30 min and stimulated with 10% fetal bovine serum (Thermo Fisher Scientific). For analysis of degradation pathways, iPSC-CMs at day 60 of differentiation were treated with 1-2 µM pevonedistat (Hycultec) or 750 ng MG-132 (InvivoGen) for three days. For each iPSC line, at least three individual differentiations/conditions were analyzed. All antibodies used for Western blot are listed in Table S2 in the supplement.

### Real-time PCR analysis of iPSC-CMs

Pellets of iPSC-CMs at day 60 of differentiation were snap-frozen in liquid nitrogen and stored at −80°C. Total RNA was isolated using the NucleoSpin RNA Mini kit (Macherey-Nagel) according to manufacturer’s instructions. 200 ng RNA was used for the first-strand cDNA synthesis by using the MULV Reverse Transcriptase and Oligo d(T)16 (Thermo Fisher Scientific). For real-time PCR, cDNA was diluted 1:1 with nuclease-free water (Promega). Quantitative real-time PCR reactions were carried out using the SYBR Green PCR master mix and ROX Passive Reference Dye (Bio-Rad) with Micro-Amp Optical 384-well plates, and the 7900HT fast real-time PCR system (Applied Biosystems) according to the manufacturer’s instructions with the following parameters: 95°C for 10 min, followed by 40 cycles at 95°C for 15 s and 60°C for 1 min. Analysis was conducted using the ΔΔCT method and values were normalized to *GAPDH* gene expression and to WT controls. Primer sequences are listed in Table S3 in the supplement.

### Analysis of sarcomere length and myofibril organization of iPSC-CMs

To analyze the sarcomere length and myofibril organization, iPSC-CMs were cultured on Matrigel-coated coverslips and fixed at day 60 of differentiation in 4% Roti-Histofix (Carl Roth) at RT for 10 min and blocked with 1% Bovine Serum Albumin (BSA; Sigma-Aldrich) in PBS (Thermo Fisher Scientific) overnight at 4°C. Primary antibodies were applied in 1% BSA and 0.1% Triton-X100 (Carl Roth) in PBS at 37°C for 1 h or at 4°C overnight. Secondary antibodies with minimal cross reactivity were administered in 1% BSA in PBS (Thermo Fisher Scientific) at RT for 1 h. Nuclei were stained with 8.1 µM Hoechst 33342 (Thermo Fisher Scientific) at RT for 10 min. Samples were mounted in Fluoromount-G (Thermo Fisher Scientific). Images were collected using the Axio Imager M2 microscopy system (Carl Zeiss) and Zen 2.3 software. For analysis of the sarcomere length, images with α-actinin staining of iPSC-CMs were evaluated using the SarcOptiM plugin in ImageJ (National Institutes of Health).^55^ Here, three independent lines along different myofibrils within one cell were selected to calculate the mean sarcomere length per cell. For each iPSC line, three individual differentiations with 9-13 images per differentiation and two cells per image were analyzed. To analyze the myofibril organization, images with α-actinin staining of iPSC-CMs were processed using the Tubeness and Fast Fourier Transform plugins in ImageJ. Processed images were radially integrated using the Radial Profile Plot plugin in ImageJ and the relative amplitude of the first-order peak in the intensity profile as a measure of sarcomere and myofibril regularity was automatically analyzed using LabChart (ADInstruments). For each iPSC line, three individual differentiations with 7-11 images per differentiation were analyzed. All antibodies used for immunofluorescence are listed in Table S2 in the supplement.

### Analysis of cell size of iPSC-CMs

To study cellular hypertrophy, iPSC-CMs at day 60 of differentiation were analyzed for cell size in suspension, as previously described.^10^ In brief, iPSC-CMs at day 50 of differentiation were plated at a density of 2.5×10^5^ cells per well on Matrigel-coated 12-well plates. At day 60 of differentiation, cells were singularized with StemPro Accutase Cell Dissociation Reagent (Thermo Fisher Scientific) and measured for cell diameter using the CASY cell counter system (OMNI Life Science). Each value represents a mean of 5×10^2^ to 1.5×10^4^ cells per measurement. To exclude cell debris and cell clusters, only values within a diameter range of 15-40 µm were selected. For each iPSC line, at least three individual differentiations with 3-5 replicates per differentiation were analyzed. To study the effect of MEK inhibition on LZTR1^L580P^ iPSC-CMs, cultured at a density of 6×10^5^ cells per well were treated with 10 nM trametinib for 5 days before being measured via the CASY cell counter.

### Video-based contractility analysis of iPSC-CMs

To analyze contractile parameters in monolayer, iPSC-CMs were cultured on Matrigel-coated 6-well plates and measured using the Cytomotion imaging setup (IonOptix). Recordings (60-75 seconds in duration) were acquired at 250 frames per second. Contractile parameters (beat frequency, beat regularity, contraction and relaxation time) were analyzed using CytoSolver.

### Ectopic expression of LZTR1 variants in iPSC-CMs

For ectopic expression studies, the human WT *LZTR1* coding sequence was synthesized (Genewiz/Azenta Life Sciences) and subcloned in *pcDNA3-HA-humanNEMO* (gift from Kunliang Guan, Addgene plasmid #13512) by exchanging the *NEMO* coding sequence. Additionally, the HA-tag was exchanged by a FLAG-tag by synthesis of the fragment and subcloning in *pcDNA3-HA-LZTR1-WT* (Genewiz/Azenta Life Sciences). Patient-specific mutations were introduced into *pcDNA3-HA-LZTR1-WT* and *pcDNA3-FLAG-LZTR1-WT* using mutagenesis PCR. Plasmid DNA was isolated via the endotoxin-free NucleoBond Xtra Midi Plus EF kit (Macherey-Nagel). For transfection, WT1 iPSC-CMs cultured on Matrigel-coated 4-well chamber slides at a density of 7×10^4^ cells per well were transfected at day 60 of differentiation with the respective plasmids using Lipofectamine Stem Transfection Reagent (Thermo Fisher Scientific) according to manufacturer’s instructions with 700 ng per plasmid. After 24 h post-transfection, cells were fixed, stained, and imaged as described above. To quantitatively analyze speckle size and filament length, a custom-build pipeline in CellProfiler (BROAD institute) was applied. For each *LZTR1* variant, plasmid transfections were performed in at least three replicates. All antibodies used for immunofluorescence are listed in Table S2 in the supplement. All plasmids used are listed in Table S4 in the supplement.

### Expression and purification of recombinant LZTR1 proteins

LZTR1^WT^ and LZTR1^L580P^ were expressed as C-terminal His-tagged proteins in Expi-293F cells (Thermo Fisher Scientific). pcDNA3.1-LZTR1-Myc-6xHis plasmid was a gift from Jens Kroll (Heidelberg University and German Cancer Research Center (DKFZ-ZMBH Alliance)).^56^ The *LZTR1^L580P^*variant was introduced into the plasmid by site-directed mutagenesis as previously described.^11^ Cells were transfected using ExpiFectamine 293 Reagent (Thermo Fisher Scientific) and cultured at a density of 3-5×10^6^ cells/ml in a 37°C incubator with ≥80% relative humidity and 8% CO2 on an orbital shaker at 125×g for 3-4 days. Expression of the recombinant LZTR1 proteins was confirmed by Western blot analysis using an anti-His tag monoclonal rabbit antibody (Thermo Fisher Scientific). Following confirmation of expression, cells were harvested and lysed in a buffer containing 50 mM Tris/HCl (pH 7.4), 250 mM NaCl, 5 mM MgCl_2_, 0.5 mM CHAPS, 0.5 mM sodium deoxycholate, 1 mM Na_3_VO_4_, 1 mM NaF, and 5% glycerol, and one complete EDTA-free protease inhibitor mixture tablet (Roche Diagnostics). The lysates were centrifuged at 20,000×g for 30 min at 4 °C to obtain the soluble protein fraction containing the expressed LZTR1 proteins. Soluble fractions were applied to a Ni-NTA resin column and bound proteins, including LZTR1 proteins, were eluted with a buffer containing 50 mM Tris/HCl (pH 7.4), 250 mM NaCl, 5 mM MgCl_2_, 5% glycerol, and 250 mM imidazole. Purified LZTR1 proteins were concentrated using a 30 kDa MWCO concentrator (Amicon), snap-frozen in liquid nitrogen, and stored at −80°C.

### Analytical size exclusion chromatography (SEC) of soluble recombinant LZTR1 proteins

Purified LZTR1 proteins were centrifuged at 12,000×g for 10 min before being applied to an analytical Superose 6 10/300 SEC column (GE Healthcare Life Sciences) using a buffer containing 50 mM Tris/HCl (pH 7.4), 250 mM NaCl, 5 mM MgCl_2_, 0.5 mM CHAPS, 0.5 mM sodium deoxycholate, and 5% glycerol at a flow rate of 0.5 ml/min. The column was calibrated using a kit (GE Healthcare Life Sciences) containing standards of known molecular weight, including blue dextran (2000 kDa), thyroglobulin (669 kDa), ferritin (440 kDa), aldolase (158 kDa), and ovalbumin (44 kDa) at their respective concentrations. The proteins were eluted with the equilibration buffer at a constant flow rate and the absorbance at 260 nm was monitored with a UV detector. The elution profiles were analyzed using OriginPro 2021 software (OriginLab) to determine the retention volume and molecular weight of the LZTR1^WT^ and LZTR1^L580P^ proteins. To ensure the accuracy of the SEC results, trichloroacetic acid precipitation of the SEC fractions was performed. The precipitated proteins were visualized by SDS-PAGE and Western blot analysis using an anti-His tag monoclonal rabbit antibody (Thermo Fisher Scientific) to determine the protein distribution in each fraction.

### Pull-down assay for analysis of LZTR1-RAS interactions

Recombinant GST-fused RAS proteins in both inactive (GDP-bound) and active (GppNHp-bound) states were prepared according to established protocols.^57^ In brief, nucleotide and protein concentrations were determined using HPLC and Bradford reagents, and aliquots were stored at −80°C. His Mag Sepharose Ni beads (GE Healthcare) were used for the protein-protein interaction assay. Recombinant LZTR1^WT^ and LZTR1^L580P^ proteins were each mixed with MRAS and RIT1 proteins in a buffer containing 50 mM Tris/HCl (pH 7.4), 250 mM NaCl, 10 mM MgCl_2_, 20 mM imidazole, and 5% glycerol. Individual protein mixtures were prepared for each LZTR1 variant and RAS protein combination. Input samples were collected for analysis, representing the initial protein composition. The remaining volume of each sample was subjected to pull-down using His Mag Sepharose Ni beads. Mixtures were incubated for 1 h at 4°C to allow for specific protein-protein interactions. After incubation, beads were thoroughly washed with binding buffer to remove non-specific binding. Protein complexes were eluted from the beads using a buffer containing 250 mM imidazole. Eluted samples were analyzed by SDS-PAGE to visualize the separated proteins. To confirm the interactions, Western blotting was performed using an anti-His tag monoclonal rabbit antibody (Thermo Fisher Scientific) and GST monoclonal mouse antibody (own antibody). GST control samples were included in each pull-down experiment to serve as negative controls, assessing the specificity of observed protein-protein interactions.

### In silico prediction of protein structures and multimer complexes

Homo-trimer configurations of the different *LZTR1* variants and configurations of LZTR1 with cullin 3 and MRAS were predicted using ColabFold (version 02c53)^30^ and AlphaFold-multimer v2^58^ with 6 recycles and no templates on an A5000 GPU with 24 GBs of RAM and repeated twice. The five predicted models for each variant were ranked according to the predicted template modeling score and interactions between the chains were inspected through the predicted alignment error generated by AlphaFold-multimer.

### Statistics

Data are presented as the mean ± standard error of the mean, unless otherwise specified. Statistical comparisons were performed using the D’Agostino-Pearson normality test and the nonparametric Kruskal-Wallis test followed by Dunn correction or the parametric t test in Prism 10 (GraphPad). Results were considered statistically significant when the p-value was ≤ 0.05.

### Data and biomaterial availability

The mass spectrometry proteomics data have been deposited to the ProteomeXchange Consortium via the PRIDE partner repository (https://www.proteomexchange.org/) with the identifiers PXD038425 and PXD038417. All human iPSC lines used in this study are deposited in the stem cell biobank of the University Medical Center Göttingen and are available for research use upon request.

## Acknowledgements

We thank Laura Cyganek, Yvonne Hintz, Nadine Gotzmann, Lisa Schreiber, and Yvonne Wedekind (Stem Cell Unit, University Medical Center Göttingen), Branimir Berečić, Tim Meyer and Malte Tiburcy (Institute of Pharmacology and Toxicology, University Medical Center Göttingen), and Anja Wiechert and Manuela Gesell Salazar (Interfaculty Institute of Genetics and Functional Genomics, University Medicine Greifswald) for excellent technical assistance.

This work was supported by the German Research Foundation (DFG): project number 417880571 to L.C.; project number 501985000 to L.C.; project number 408077919 to I.C.C.; project number 193793266, Collaborative Research Centre 1002, C04, D01, D02 and S01 to W.H.Z., G.H., B.W. and L.C.; project number 390729940, Germany’s Excellence Strategy - EXC 2067/1 to A.V.B., G.H., W.H.Z., B.W. and L.C.; by the Else Kröner Fresenius Foundation: project number 2019_A75 to L.C.; by the German Federal Ministry of Education and Research (BMBF)/German Center for Cardiovascular Research (DZHK) to E.H., G.H., W.H.Z., B.W. and L.C.; by the German Federal Ministry of Education and Research (BMBF): GeNeRARe project numbers 01GM1902F and 01GM1519D, to I.C.C. and R.A.; and by the Leducq Foundation: project number 20CVD04 to W.H.Z.

## Author contributions

L.C. designed the study. L.C. and A.V.B. designed the experiments. A.V.B., O.G.G., E.H., F.K., A.M., M.S., C.P., L.B., H.S., M.K., M.E., J.A., F.M., and L.C. performed the experiments and analyzed the data. I.C.C., G.H., W.H.Z., R.A., and B.W. gave technical support and conceptual advice. L.C. and A.V.B. wrote and edited the manuscript.

---

**Figure S1:**
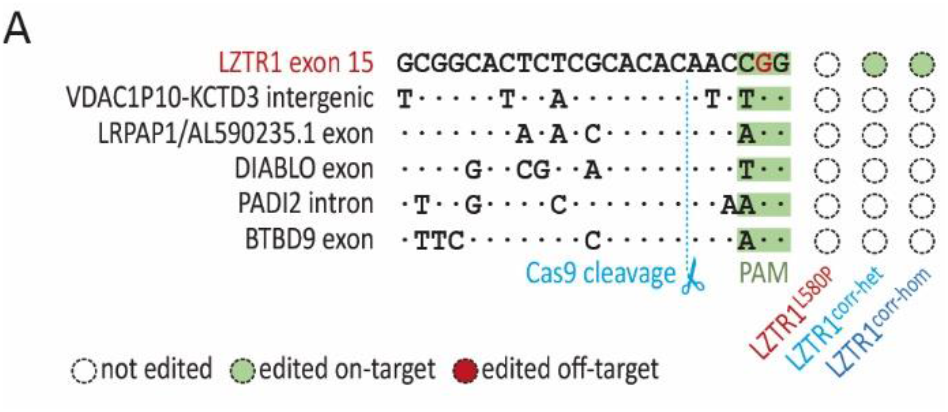
Off-target screening in CRISPR/Cas9-edited iPSCs. **(A)** Sanger sequencing of the top five predicted off-target regions, ranked by the CFD off-target score using CRISPOR, revealed no off-target editing of CRISPR/Cas9 in CRISPR-corrected iPSCs compared to the patient-derived cells.

**Figure S2:**
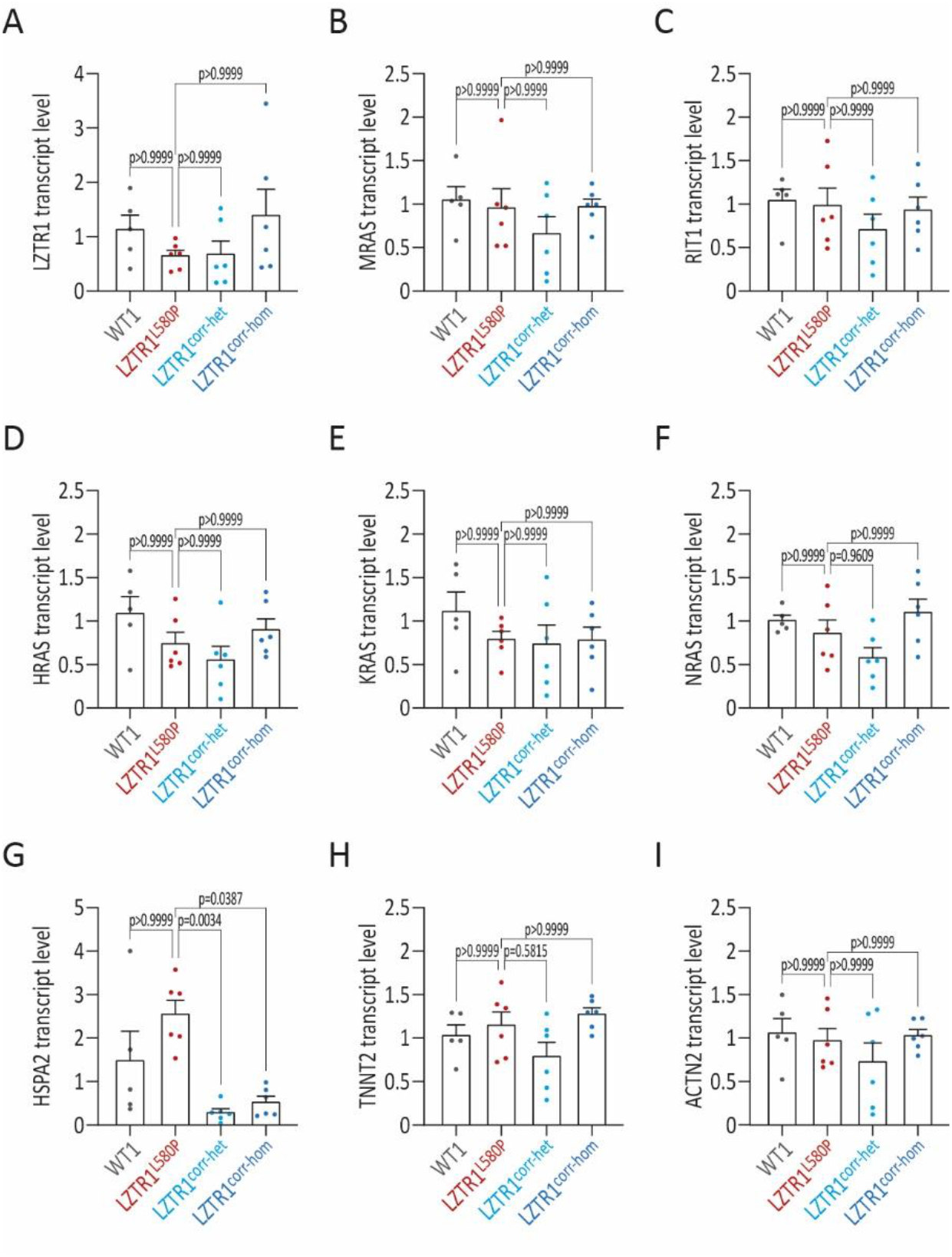
Homozygous *LZTR1^L580P^*shows no upregulation of RAS GTPases at transcriptional level. **(A-I)** Quantitative gene expression analysis of *LZTR1* (A), of LZTR1 substrates *MRAS* (B), *RIT1* (C), *HRAS* (D), *KRAS* (E), and *NRAS* (F), of *HSPA2* (G), and of cardiac-specific genes *TNNT2* (H), and *ACTN2* (I) in WT, the patient-specific, and the two CRISPR-corrected iPSC-CMs at day 60 of differentiation, assessed by real-time polymerase chain reaction, revealed no expression differences at transcriptional level across all iPSC lines; samples were analyzed in duplicates and data were normalized to *GAPDH* expression and WT controls; n=5-6 independent differentiations per iPSC line. Data were analyzed by nonparametric Kruskal-Wallis test with Dunn correction and are presented as mean ± SEM (A-I).

**Figure S3:**
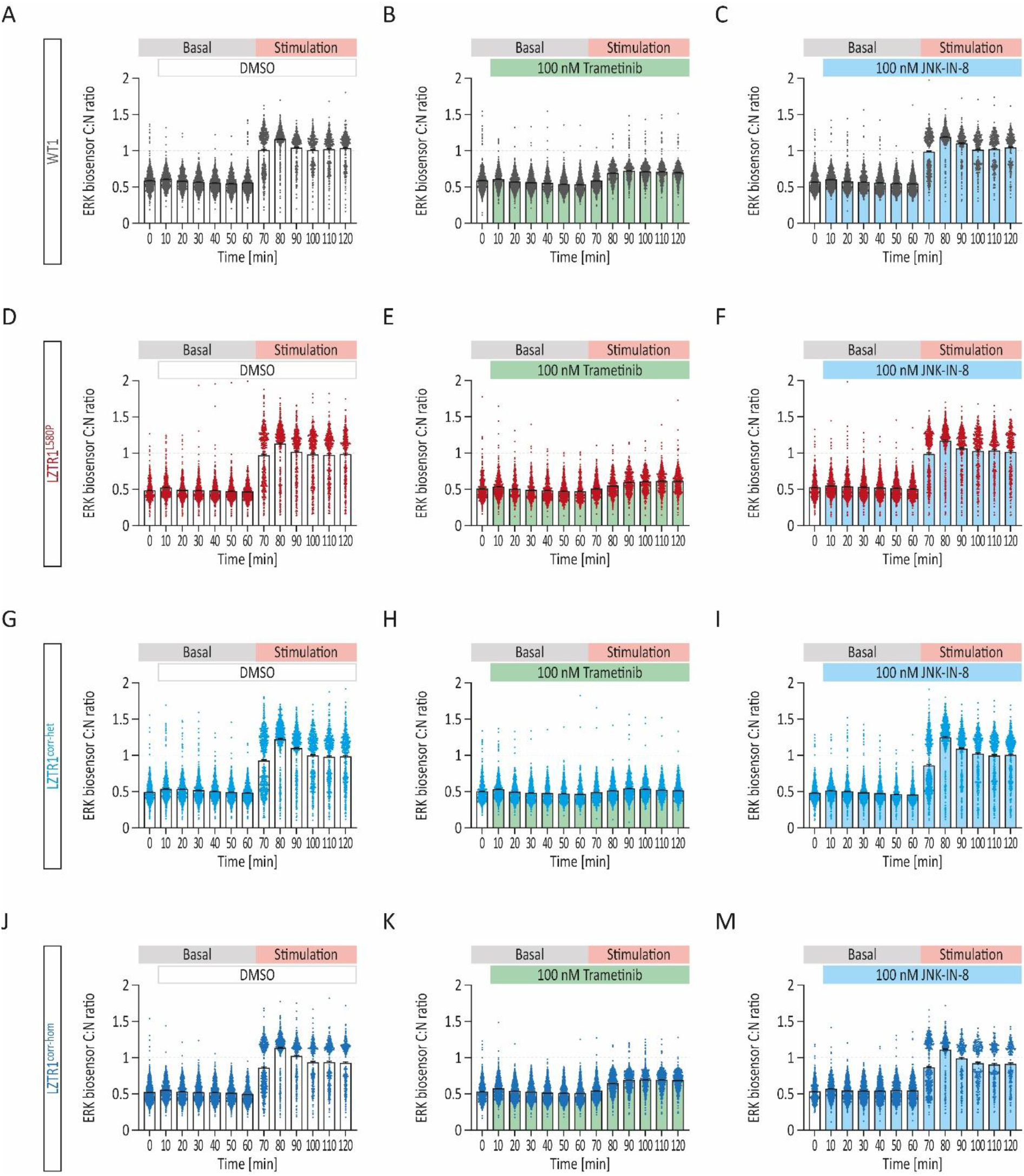
Biosensor-based analysis of ERK signaling dynamics in real time. **(A-M)** Quantitative analysis of ERK biosensor cytosol/nucleus (C:N) ratio in WT (A-C), the patient-specific (D-F), and the two CRISPR-corrected (G-M) biosensor-transduced iPSC-CMs treated with MEK inhibitor trametinib, with JNK inhibitor JNK-IN-8, or with DMSO for 60 minutes, before stimulation with serum for another 60 minutes; n=2 independent differentiations per iPSC line with n=4-5 individual wells per condition.

**Figure S4:**
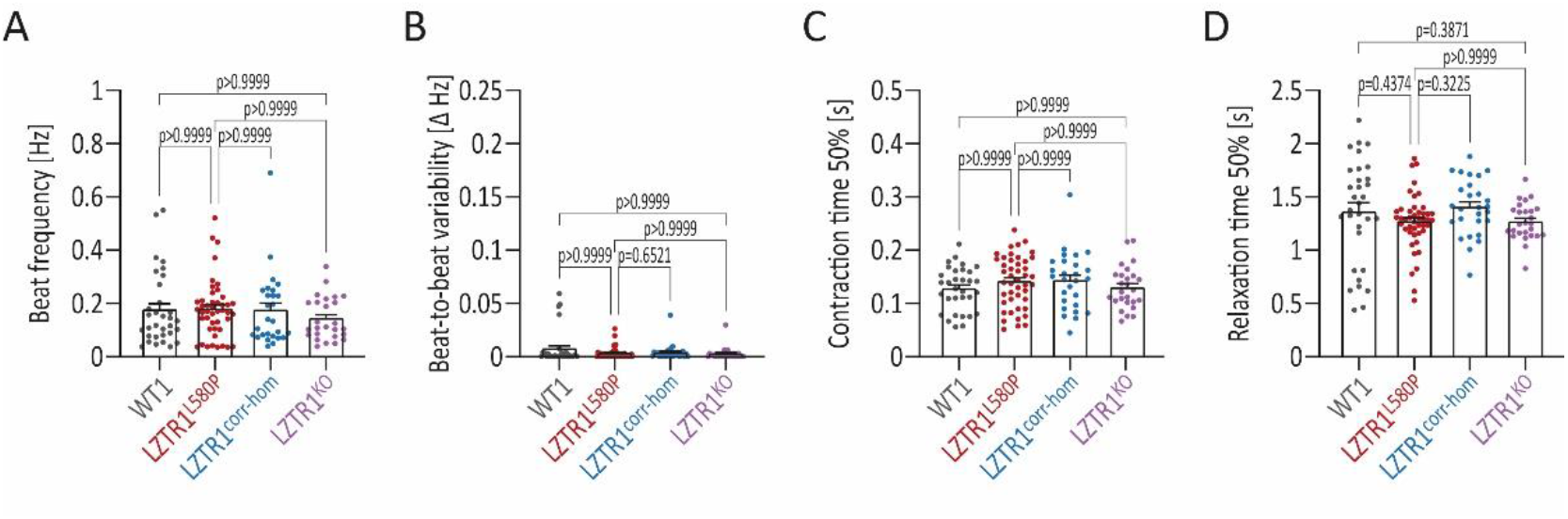
Homozygous *LZTR1^L580P^*shows unchanged contractile properties. **(A-D)** Quantitative analysis of beating frequency (A), beat-to-beat variability (B), contraction time (C), and relaxation time (D) in WT, the patient-specific, the homozygous CRISPR-corrected, and LZTR1^KO^ iPSC-CMs at day 60 of differentiation, assessed by video-based contractility analysis in monolayer cultures, revealed no significant differences in contractile function across all iPSC lines; n=3-9 independent differentiations per iPSC line. Data were analyzed by nonparametric Kruskal-Wallis test with Dunn correction and are presented as mean ± SEM (A-D).

**Figure S5:**
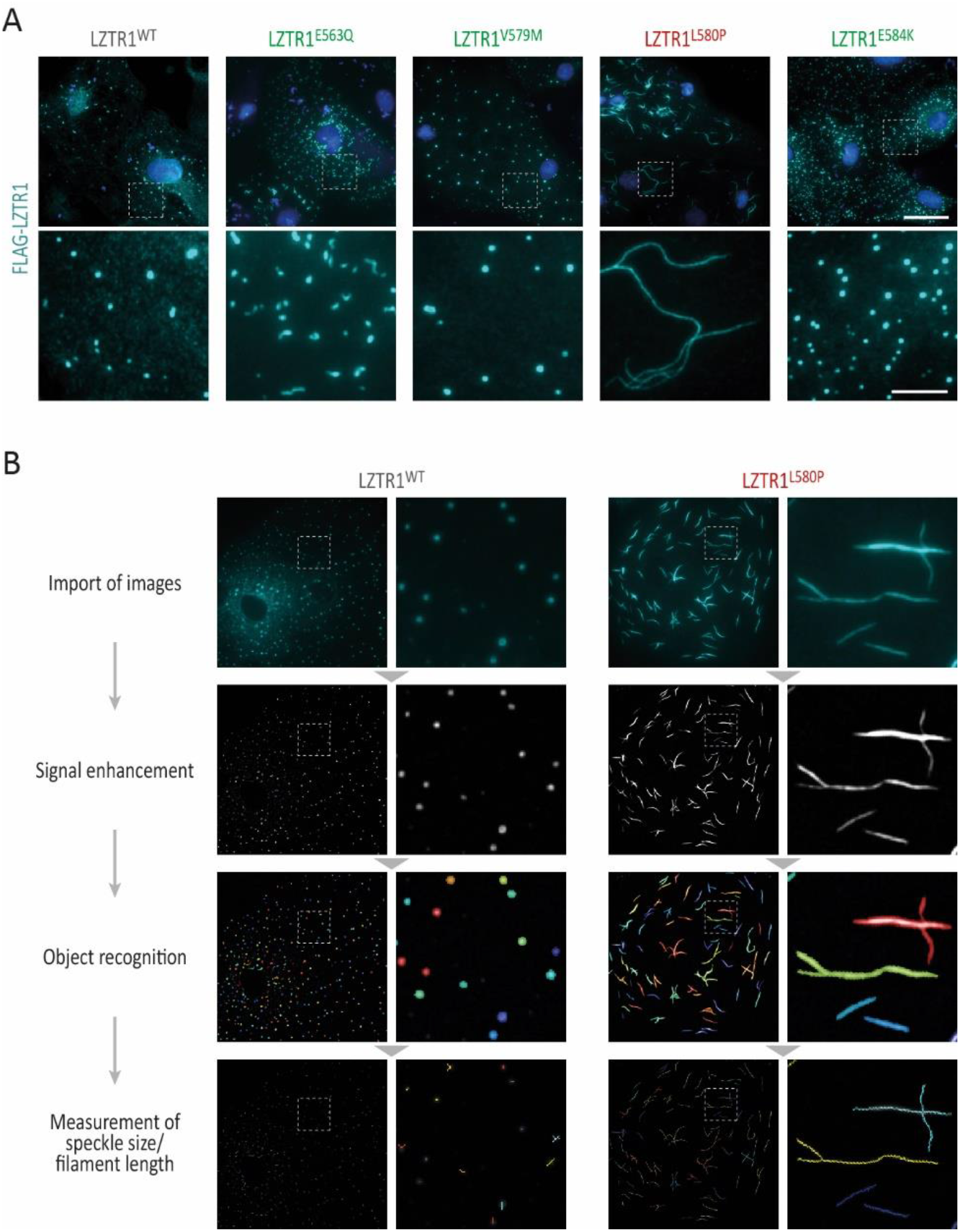
Unique *LZTR1^L580P^*-induced polymerization of LZTR1 complexes. **(A)** Representative images of WT iPSC-CMs at day 60 of differentiation after single plasmid transfection stained for FLAG-tagged LZTR1 confirmed that only LZTR1^L580P^ forms large filaments, whereas LZTR1^WT^ and the other variants present a speckle-like pattern; nuclei were counter-stained with Hoechst 33342 (blue); scale bars: 20 µm in upper panel, 5 µm in lower panel. **(B)** Customized CellProfiler pipeline for recognition and quantification of speckle size and filament length in iPSC-CMs with ectopic expression of LZTR1 variants.

**Figure S6:**
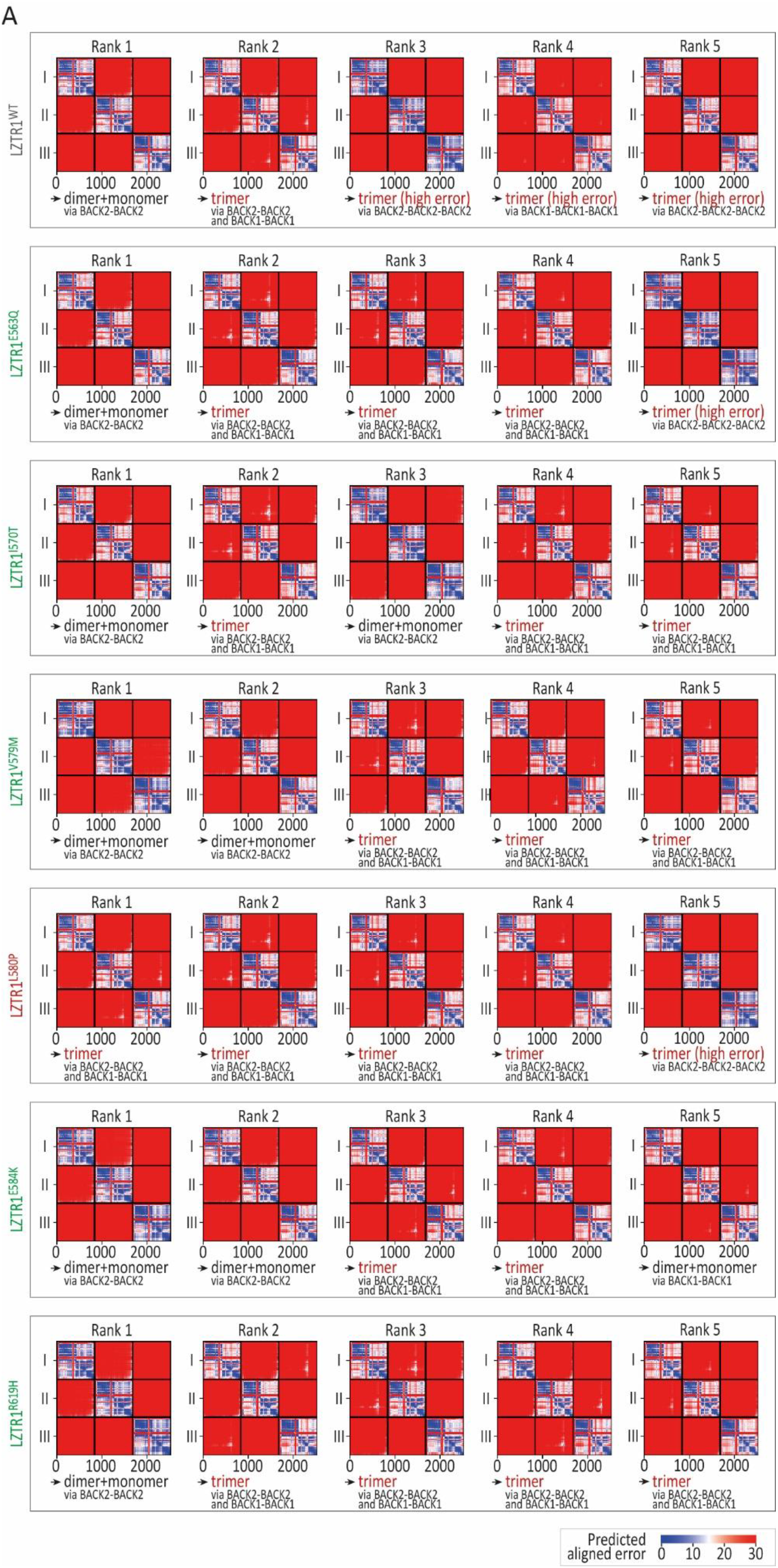
Computational prediction for LZTR1 interactions via ColabFold. (A) The five predicted models for each LZTR1 variant were ranked according to the predicted template modeling score and interactions between the chains were inspected through the predicted alignment error generated by AlphaFold-multimer.

**Table S1:** Clinical characterization of the affected patient.

|  |
| --- |
| **Cardiac findings** |
| Mild left ventricular hypertrophy |
| Prolonged QT interval |
| Stress-induced cardiac arrhythmias |
| Pericardial effusion |
| **Facial characteristics** |
| Down-slanting palpebral fissures |
| Mild bilateral ptosis |
| Triangular facial contour |
| Curly hair |
| Low posterior hairline |
| High-arched palate |
| **Physical characteristics** |
| Marfanoid habitus:<br>height 186 cm (75th-90th percentile)<br>weight 56 kg (3th percentile) |
| Pronounced pectus excavatum |
| Scoliosis |
| Stretch marks on lower back |
| Clinodactyly |
| **Additional findings** |
| Mild bilateral sensorineural hearing loss |

**Table S2:**
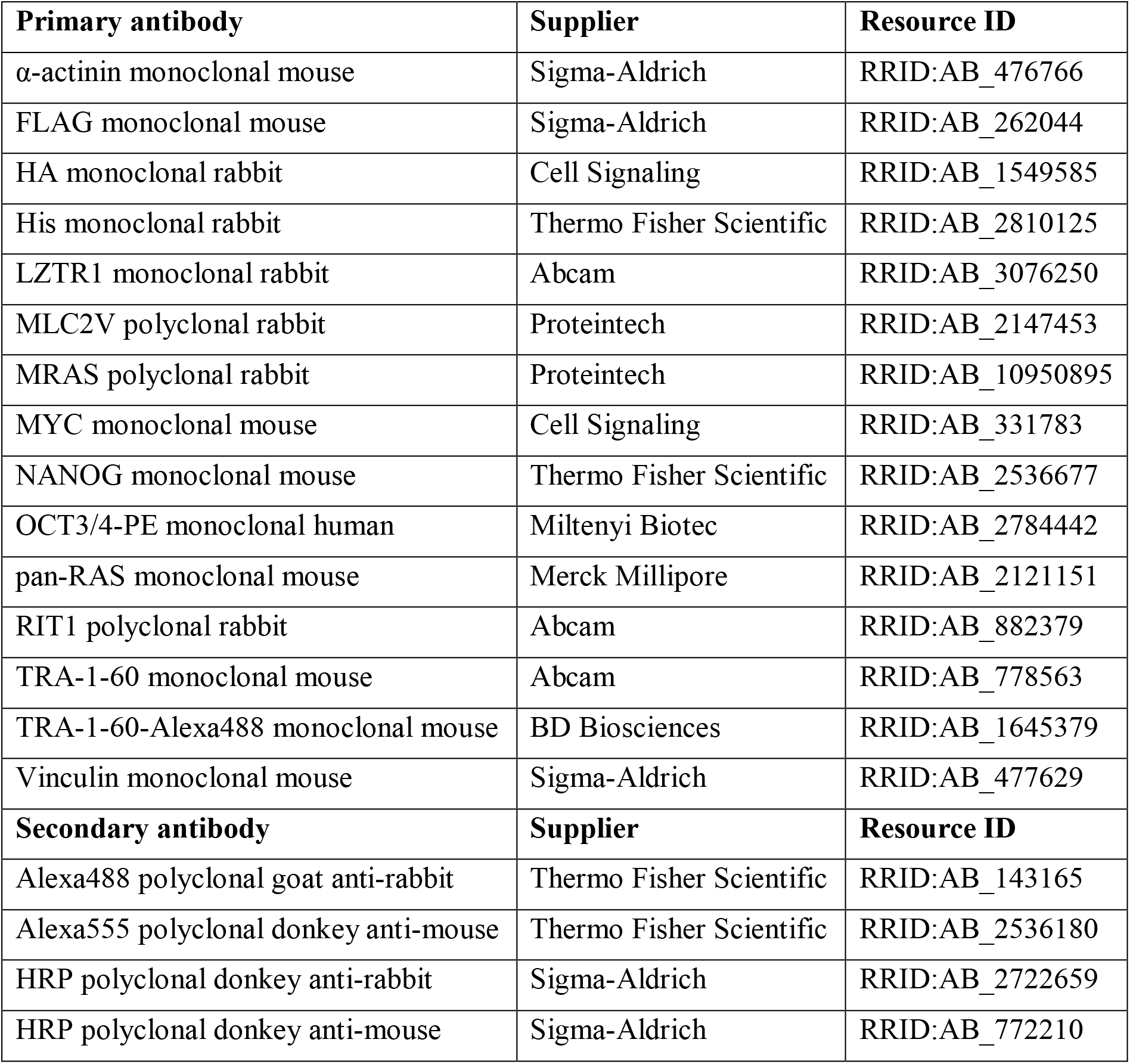
Antibodies used for Western blot, immunocytochemistry and flow cytometry.

| **Primary antibody** | **Supplier** | **Resource ID** |
| --- | --- | --- |
| α-actinin monoclonal mouse | Sigma-Aldrich | RRID:AB_476766 |
| FLAG monoclonal mouse | Sigma-Aldrich | RRID:AB_262044 |
| HA monoclonal rabbit | Cell Signaling | RRID:AB_1549585 |
| His monoclonal rabbit | Thermo Fisher Scientific | RRID:AB_2810125 |
| LZTR1 monoclonal rabbit | Abcam | RRID:AB_3076250 |
| MLC2V polyclonal rabbit | Proteintech | RRID:AB_2147453 |
| MRAS polyclonal rabbit | Proteintech | RRID:AB_10950895 |
| MYC monoclonal mouse | Cell Signaling | RRID:AB_331783 |
| NANOG monoclonal mouse | Thermo Fisher Scientific | RRID:AB_2536677 |
| OCT3/4-PE monoclonal human | Miltenyi Biotec | RRID:AB_2784442 |
| pan-RAS monoclonal mouse | Merck Millipore | RRID:AB_2121151 |
| RIT1 polyclonal rabbit | Abcam | RRID:AB_882379 |
| TRA-1-60 monoclonal mouse | Abcam | RRID:AB_778563 |
| TRA-1-60-Alexa488 monoclonal mouse | BD Biosciences | RRID:AB_1645379 |
| Vinculin monoclonal mouse | Sigma-Aldrich | RRID:AB_477629 |
| **Secondary antibody** | **Supplier** | **Resource ID** |
| Alexa488 polyclonal goat anti-rabbit | Thermo Fisher Scientific | RRID:AB_143165 |
| Alexa555 polyclonal donkey anti-mouse | Thermo Fisher Scientific | RRID:AB_2536180 |
| HRP polyclonal donkey anti-rabbit | Sigma-Aldrich | RRID:AB_2722659 |
| HRP polyclonal donkey anti-mouse | Sigma-Aldrich | RRID:AB_772210 |

**Table S3:**
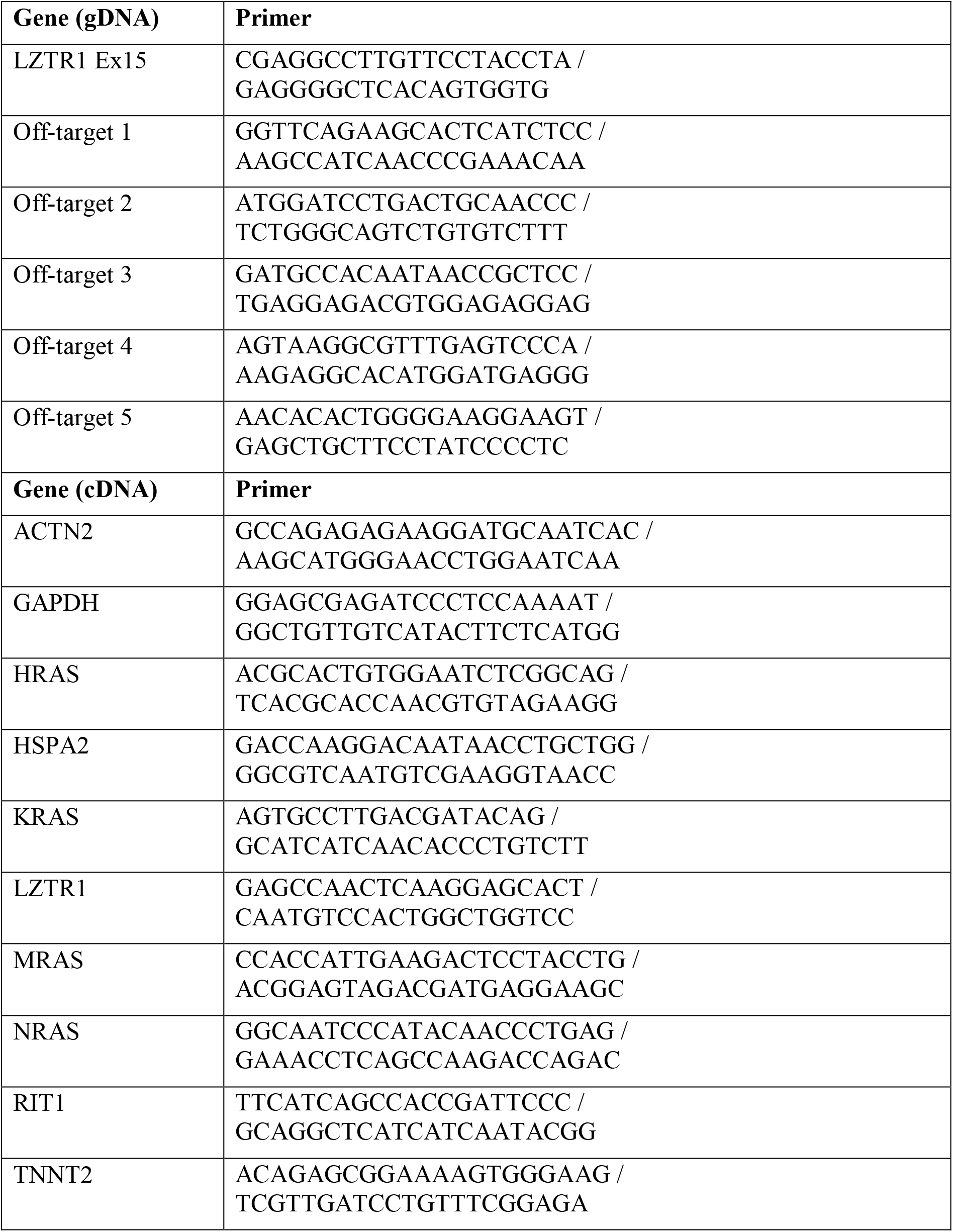
Primer sequences used for PCR and real-time PCR.

**Table S4:** Plasmids used in this study.

| Plasmid | Source |
| --- | --- |
| pcDNA3-HA-LZTR1-WT | modified from RRID:Addgene_13512 |
| pcDNA3-FLAG-LZTR1-WT | modified from pcDNA3-HA-LZTR1-WT |
| pcDNA3-HA-LZTR1-E563Q | modified from pcDNA3-HA-LZTR1-WT |
| pcDNA3-FLAG-LZTR1-E563Q | modified from pcDNA3-FLAG-LZTR1-WT |
| pcDNA3-HA-LZTR1-I570T | modified from pcDNA3-HA-LZTR1-WT |
| pcDNA3-HA-LZTR1-V579M | modified from pcDNA3-HA-LZTR1-WT |
| pcDNA3-FLAG-LZTR1-V579M | modified from pcDNA3-FLAG-LZTR1-WT |
| pcDNA3-HA-LZTR1-L580P | modified from pcDNA3-HA-LZTR1-WT |
| pcDNA3-FLAG-LZTR1- L580P | modified from pcDNA3-FLAG-LZTR1-WT |
| pcDNA3-HA-LZTR1-E584K | modified from pcDNA3-HA-LZTR1-WT |
| pcDNA3-FLAG-LZTR1-E584K | modified from pcDNA3-FLAG-LZTR1-WT |
| pcDNA3-HA-LZTR1-R619H | modified from pcDNA3-HA-LZTR1-WT |
| pcDNA3-HA-LZTR1-ΔBTB2-BACK2 | modified from pcDNA3-HA-LZTR1-WT |
| pLentiPGK Puro DEST ERKKTRClover | RRID:Addgene_90227 |
| pMD2.G | RRID:Addgene_12259 |
| psPAX2 | RRID:Addgene_12260 |
| pcDNA3.1-LZTR1-Myc-6xHis | Jens Kroll (Heidelberg University and German Cancer Research Center)<sup>56</sup> |
| pcDNA3.1-LZTR1-L580P-Myc-6xHis | modified from pcDNA3.1-LZTR1-Myc-6xHis |

